# The common message of constraint-based optimization approaches: overflow metabolism is caused by two growth-limiting constraints

**DOI:** 10.1101/679019

**Authors:** Daan H. de Groot, Julia Lischke, Riccardo Muolo, Robert Planqué, Frank J. Bruggeman, Bas Teusink

## Abstract

Living cells can express different metabolic pathways that support growth. The criteria that determine which pathways are selected in which environment remain unclear. One recurrent selection is overflow metabolism: the simultaneous usage of an ATP-efficient and -inefficient pathway, shown for example in *Escherichia coli*, *Saccharomyces cerevisiae* and cancer cells. Many models, based on different assumptions, can reproduce this observation. Therefore, they provide no conclusive evidence which mechanism is causing overflow metabolism. We compare the mathematical structure of these models. Although ranging from Flux Balance Analyses to self-fabricating Metabolism and Expression models, we can rewrite all models into one standard form. We conclude that all models predict overflow metabolism when two, model-specific, growth-limiting constraints are hit. This is consistent with recent theory. Thus, identifying these two constraints is essential for understanding overflow metabolism. We list all imposed constraints by these models, so that they can hopefully be tested in future experiments.

## 1 Introduction

Many cells show overflow metabolism: the simultaneous metabolism of nutrients by an energy-efficient and a less energy-efficient pathway. For example, *Escherichia coli*, *Saccharomyces cerevisiae* and cancer cells fully oxidize carbon sources to CO_2_ when growing slowly. Above a species-specific critical growth rate, a partial oxidation pathway kicks in, resulting in the production of overflow metabolites: acetate, ethanol and lactate respectively [1–3]. *Lactococcus lactis* shows a similar metabolic shift from mixed-acid fermentation (3 ATP per glucose) to lactic-acid fermentation (2 ATP per glucose) under anaerobic conditions [4]. Besides overflow metabolism that starts at high growth rates, *Escherichia coli* even produces overflow products at low growth rates when growing in ammonium-limited conditions [5].

Overflow metabolism seems wasteful because two metabolic pathways are used that independently support growth, and one of them is more efficient (it has a higher ATP yield per glucose molecule) than the other. Since cells need energy for growth, efficient usage of nutrients is expected to be favourable. One would therefore expect that cells using the efficient growth strategy exclusively would be selected during evolution.

The counterintuitive occurrence of overflow metabolism is in many studies explained using constraint-based optimization approaches. These approaches assume that cellular growth is constrained by physical and chemical limits, and that cells are driven towards these limits when evolutionary fitness is maximized. Accordingly, the behavior of cells results from maximizing their growth rate given a set of constraints.

Since many models reproduce the experimental data while using different biological assumptions, it is unclear what exactly causes overflow metabolism. Therefore, we need a way to analyze and compare these different models to find the cause of overflow metabolism.

Minimal, growth-supporting metabolic modes are characterized mathematically by identifying the smallest subnetworks of the entire metabolic network that can support growth. Such subnetworks are called Elementary Flux Modes (EFMs) in metabolic models [6], and Elementary Growth Modes (EGMs) in self-fabrication models [7] (see SI1 for a short introduction and comparison of EFMs and EGMs). The gradual transition from the usage of one metabolic subnetwork to the mixed usage of two subnetworks that is observed in overflow metabolism indicates the simultaneous usage of two different Elementary Modes.

In recent theoretical work [7,8] we derived that cells that maximize their growth rate will only use two Elementary Modes if they are confronted with at least two constraints. The identification of these constraints is therefore an important step towards finding the mechanistic cause of overflow metabolism. In this review, we use this theory to compare the various models of overflow metabolism by making the growth-limiting constraints explicit.

Although the models range from relatively simple Flux Balance Analyses to genome-scale self-fabrication models, we will show that they can all be written in the same concise standard form. Thus, the models are highly similar: (a proxy for) the cellular growth rate is maximized subject to two constraints. However, the biological assumptions underlying the imposed constraints differ between those models. Hence, the success of these models is dependent on the existence of two constraints and not on the precise biological interpretation of those constraints. Finding the causes of overflow metabolism therefore amounts to identifying the two active growth-limiting constraints and experimentally testing them. We shall conclude that the models each offer a hypothesis that needs to be tested in falsification experiments in the future.

## 2 A standard form for overflow metabolism models

We will show that, to our knowledge, all existing models that use growth rate maximization to explain overflow metabolism, can be rewritten in a standard form.

We will assume that a cell adapts its state to grow as fast as possible whenever it encounters a new environment. The cellular state is specified by *optimization variables*, for example the reaction rates (***v***) or the enzyme concentrations (***e***). We will denote the optimization variables by the vector ***x***, the *i*^th^ entry of which is denoted by *x*_*i*_. The growth rate is modeled as a linear function: *the objective function*:

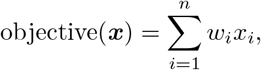

where *w*_*i*_ is the weighting factor of variable *i*. The growth rate maximization of the cell is modeled mathematically by searching for the set of optimization variables that maximizes the objective function, given constraints to be specified later. Because the objective function is linear, there is a certain direction in the space of optimization variables in which the objective always increases. The optimal solution is the set of optimization variables that is as far in that direction as possible.

Not all combinations of optimization variables can be chosen due to *constraints*, for example a limited uptake rate, or a limited available area for membrane proteins. These constraints are formalized by inequalities acting on a weighted sum of the variables:

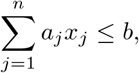

where *a*_*j*_ determines the ‘cost’ of increasing the *j*^th^ variable. In the special case that *a*_*j*_ = 0, *x*_*j*_ is not bounded by this constraint. In general, we could have several, say *m*, constraints. These constraints can be collected in an *m*×*n* matrix *A*, where the *i*^th^ row captures the *i*^th^ constraint. All constraints can then be written together as:

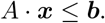

The constraints can be viewed as planes bounding the feasible combinations of variables (see Figure 1). After all constraints have been implemented, we are left with an angular space called *the solution space*. Solving the optimization problem amounts to selecting the point in this space that maximizes the objective function. It can be shown that there is always a corner point of the solution space (called vertex) in which this optimum is attained.

**Fig. 1.**
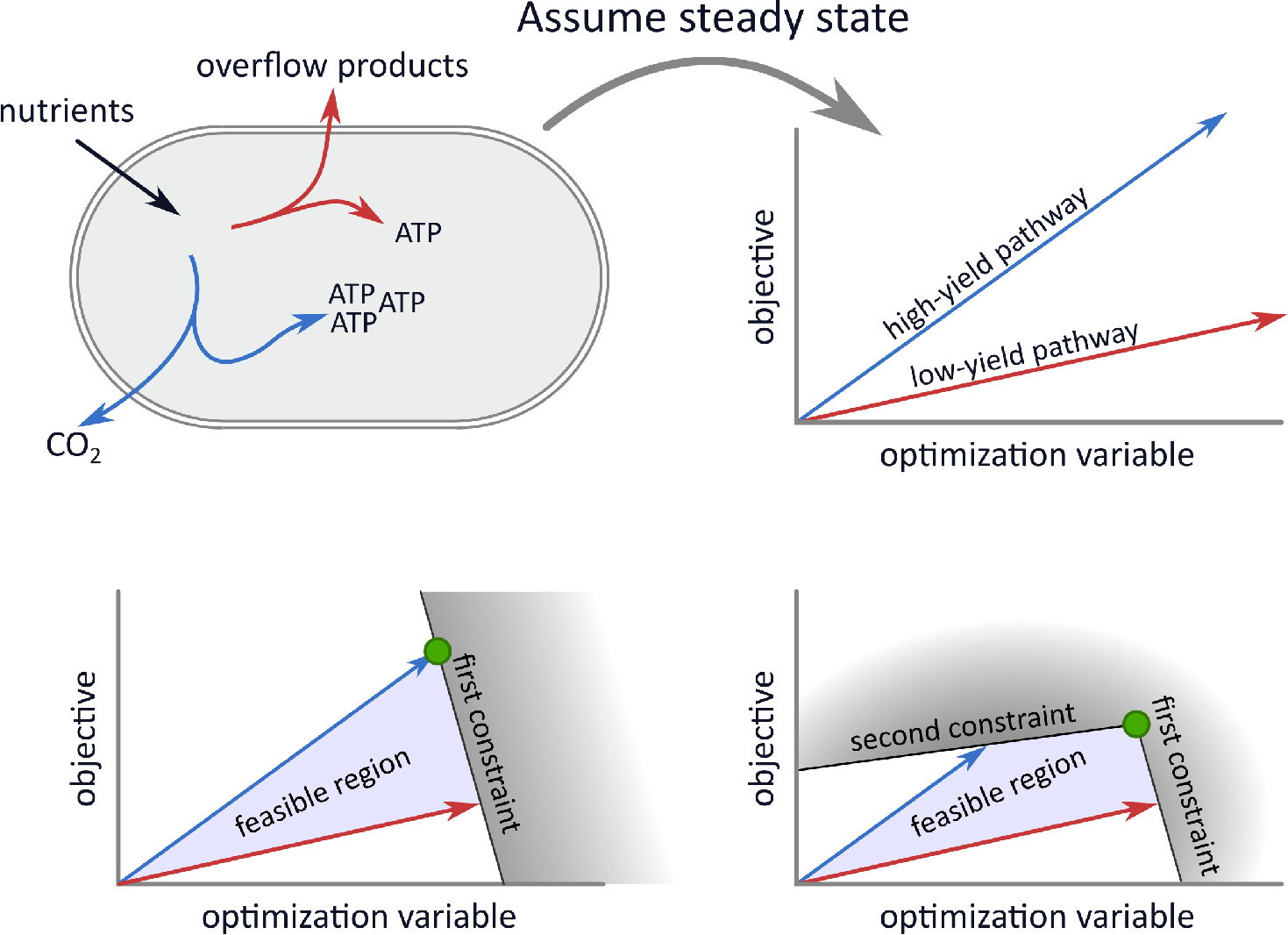
A general view on overflow metabolism and how it is modeled. In general, overflow metabolism is the simultaneous usage of two independent growth-supporting subnetworks with different substrate yields. In the **top left** subfigure, the blue pathway produces more energy equivalents per gram nutrient than the red pathway. Together with the non-depicted rest of the metabolic network, the blue and red pathway can separately lead to steady state growth. In the **top right** subfigure, we illustrate that imposing *homogeneous constraints*, in this case a steady state assumption, gives rise to relations between optimization variables. The optimization variables can for example be reaction rates or enzyme concentrations, but for simplicity, we only show one variable here. The model objective is here visualized along the *y*-axis, so that the combination of variables that gives the highest *y*-coordinate is optimal. In the **bottom** figures, we add *inhomogeneous constraints* on the optimization variables. These affect which combination of variables is optimal. Under one constraint, exclusive usage of the high-yield pathway is optimal. Adding the second constraint leads to the optimality of a combination of the two pathways.

In this review, we will use these concepts to extract the mathematical cores of all overflow metabolism models (that we could find) and rewrite them in the following standard form.^1^

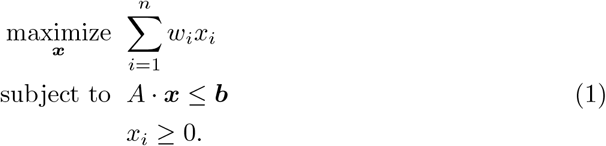

The constraints in *A* can be *homogeneous* and *inhomogeneous*, see Figure 1 and its caption. A constraint is called homogeneous when the corresponding weighted sum of the variables equals zero, i.e., *b* = 0, and inhomogeneous otherwise. Examples of homogeneous constraints that we will define later in this review are the steady state constraint and irreversibility constraints. When all constraints of *A* are homogeneous, the solution space is unbounded; it can be visualized as an infinitely stretched angular cone. The optimization problem will in this case not have a finite maximum. Inhomogeneous constraints can make the cone bounded; this will be especially important in modeling overflow metabolism. We will therefore highlight them by presenting them separate from the rest of the constraints.

## 3 Current explanations of mixed behaviour and their mathematical background

Next, we will discuss published models made to explain overflow metabolism that use growth rate maximization. We will start with the modeling approaches that are the easiest to understand, and gradually build up complexity, ending with self-fabrication models.

### 3.1 Flux Balance Analysis models

Flux Balance Analysis studies the sets of reaction rates (fluxes) through a metabolic network (say with *m* metabolites and *r* reactions) that can reach a *steady state*. A steady state is attained when the net rate of production of each metabolite is equal to the net rate of its consumption. The stoichiometry of all reactions is described by the stoichiometric matrix, *N*, which has *m* rows and *r* columns. Each row corresponds to the mass balance of a metabolite and contains the stoichiometric coefficients of this metabolite in all reactions. When *N* is multiplied by the rate vector, we get the ‘massbalance constraints’:

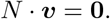

In this review we will consider all reactions to be irreversible; we can always split up a reversible reaction into one forward and one backward reaction, resulting in:

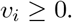

These steady state and irreversibility constraints are the homogeneous constraints. As mentioned before, the space of flux vectors that satisfy these constraints is unbounded.

In addition, flux bounds can be imposed. Upper bounds, denoted *ub*_*i*_, are for example imposed to model a limited capacity of the cell for the corresponding reaction. Lower bounds, denoted *lb*_*i*_, are for instance used to model the production of ATP for non-growth associated maintenance. This gives

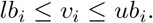

In FBA, we are mostly interested in those flux vectors, ***v***, that maximize some proxy for the growth rate. For this, the so-called *biomass reaction*, *v*_*BM*_ [9], is added to the model: a phenomenological reaction that produces all cellular compounds in the right proportions, and thereby approximates the demands for cell synthesis.

The full problem can now be written as

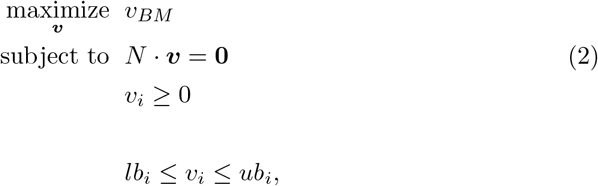

which is equivalent to the standard form that we introduced in Equation (1) (see the Supporting Information for the appropriate choice of ***w***, *A*, ***x***, ***b***).

FBA models have been used to explain overflow metabolism, mathematically capturing the reasoning of Andersen and von Meyenburg [10] in 1980: “If, however, respiration is limited, by-product formation can lead to extra ATP production and to faster growth, provided the by-product can be generated with a net gain of ATP.” The imposed flux bounds differ between the models, although all models consider a limited uptake rate for the carbon source. For example, Majewski and Domach [11] further propose that *E. coli* might have a limited electron transfer capacity, while Varma and Palsson [12] assume that oxygen uptake is limited, and that a certain amount of ATP should be produced even if the cell is not growing. This leads to the following FBA problem:

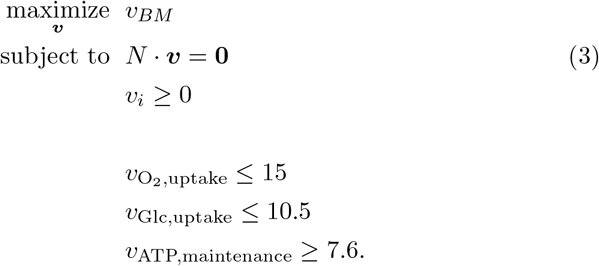

Carlson and Srienc [13,14] also model growth rate maximization under glucose- and oxygen-limitation, but take a different approach. Instead of finding only the optimal solution, they characterize the whole steady-state solution space by enumerating the EFMs of a coarse-grained *E. coli* network (see SI1 for an explanation about EFMs). Using their acquired knowledge of all possible solutions, the authors select four Elementary Flux Modes. Under any level of glucose and oxygen limitation, two of these EFMs together form the optimal solution. The simultaneous usage of these EFMs leads to overflow metabolism.

### 3.2 FBA models with thermodynamic constraints

FBA models can be refined by adding thermodynamic constraints [15, 16]. The laws of thermodynamics dictate that a chemical reaction can only have a positive rate if the summed Gibbs free energy of the reaction substrates is higher than of the reaction products, i.e., if the free energy change due to the reaction is negative, denoted by: *Δ*_*r*_*G*′ < 0. By using this, one for example excludes cycles like *A* → *B* → *C* → *A* from carrying a positive flux, since such a cycle has zero thermodynamic driving force [17]. The free energy change due to a reaction depends on the concentrations of the involved metabolites, but these are usually not modeled in FBA approaches. Most thermodynamic FBA approaches thus need some way to estimate either these metabolite concentrations, or the *Δ*_*r*_*G*′-values directly. There are also some methods where this estimation step can be avoided, at the expense of the thermodynamic constraints becoming less restrictive (see [16] for an overview of thermodynamic FBA methods).

Recently, Niebel et al. combined growth rate maximization and a thermodynamic constraint to describe overflow metabolism [18]. In their approach, the metabolite concentrations and the reaction rates are free variables, although the metabolite concentrations are provided with an upper and lower bound based on experimental measurements. The authors search for the optimal concentrations and rates so that the biomass production rate is maximized. This search is constrained by the second law of thermodynamics, implying that the free energy change induced by an active reaction should be negative: 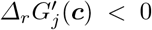 for reactions with *v*_*j*_ > 0. These homogeneous constraints take the place of the irreversibility constraints that were used in FBA models, where the directionality is now based on the sign of the Gibbs free energy change.

If we add up all these free energy changes induced by the chemical reactions, we get the total dissipated Gibbs energy per unit time: 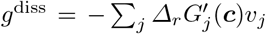. The authors observed in experiments that this dissipation function appears to have a maximum at the onset of overflow metabolism. Therefore, they propose that the dissipated energy might be limited by an upper bound,

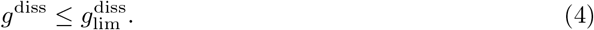

In addition, another constraint is imposed ensuring that the free energy dissipated by internal reactions equals the free energy that is extracted from external nutrients. However, after a careful examination of the mathematics used in [18] (see SI4), we believe that this constraint should be equivalent to the steady state assumption, so that we could ignore it here. If it turns out that we are wrong, the constraint can be added to the problem below, without affecting the conclusions of this review.

This modeling approach is no longer linear in the variables because the 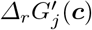-values can depend non-linearly on the metabolite concentrations. However, for any fixed set of metabolite concentrations, ***c*** = ***c***_0_, the model reduces to a Linear Program that can be written in our standard form (see SI3.2 for the appropriate choice of the variables ***w***; ***A***; ***x***; ***b*** used in (1)).

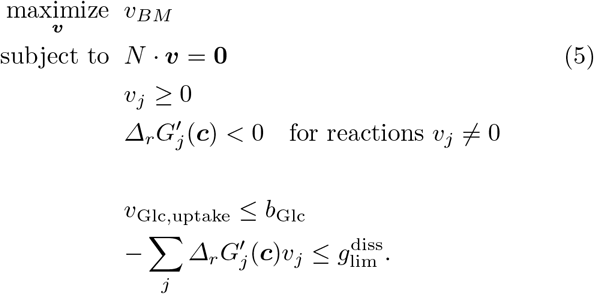

### 3.3 Resource-allocation models

Reaction rates can almost always be increased by increasing the concentration of the catalyzing enzyme [19]. A constraint on a reaction rate can therefore not reflect the mechanistic cause of metabolic phenomena: if a cell would be confronted with such a constraint, the concentration of the corresponding enzyme could be increased,unless the enzyme concentration itself is constrained. In that case however, it is the constraint on enzyme concentrations that is the mechanistic cause.

In the past decade, many researchers shifted perspective by taking enzyme concentrations as the optimization variables instead of the reaction rates. These models are called *resource allocation* models [20–31].

Resource allocation models also maximize the biomass reaction rate, *v*_BM_, while metabolism is at steady state: *N* · ***v*** = **0**. The rate of the objective reaction can thus only be increased if the rates of all reactions in a complete growth-supporting subnetwork are increased. Unlike in FBA models however, each reaction rate is now coupled to the concentration of a catalyzing enzyme,

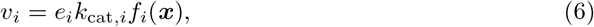

where *v*_*i*_ is the *i*^th^ reaction rate and *e*_*i*_ is the concentration of the corresponding enzyme. The activity of an enzyme is determined by its catalytic rate *k*_cat,*i*_, and the ‘saturation’ of the enzyme *f*_*i*_(***x***) with its substrates ***x***. This saturation term is in reality a nonlinear function of the metabolite concentrations ***x***, that also includes product inhibition. However, we split reversible reactions, product inhibition is almost always ignored, and *f*_*i*_(***x***) is often simplified to be constant, such that *v*_*i*_ = *e*_*i*_*k*_cat,*i*_ where *k*_cat,*i*_ is now an effective rate constant. The only way to increase the reaction rates is then to increase the enzyme concentrations. However, resources are limited: various limits on enzyme concentrations exist, which take the form of (weighted) sums that are bounded:

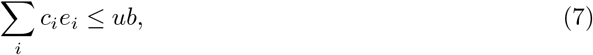

where *c*_*i*_ is a weighting factor, and should not be confused with the metabolite concentrations that were used earlier. All enzymes for which the weighting factor is nonzero, *c*_*i*_ > 0, contribute to the sum. These weighting factors can be adjusted to capture various constraints. For example, if the membrane area is constrained, the weight *c*_*i*_ would reflect the area taken up by one unit of protein *i*, and *c*_*i*_ would thus be zero for all non-membrane proteins. Since the sum is bounded, an increase in the concentration of protein *i* must be compensated by a decrease in the concentration of others. The available resources should thus be carefully allocated in order to maximize the biomass production rate. These approaches can be written in a form equivalent to our standard form (see SI3.3 for the appropriate choice of ***w***, *A*, ***x***, ***b*** in (1))

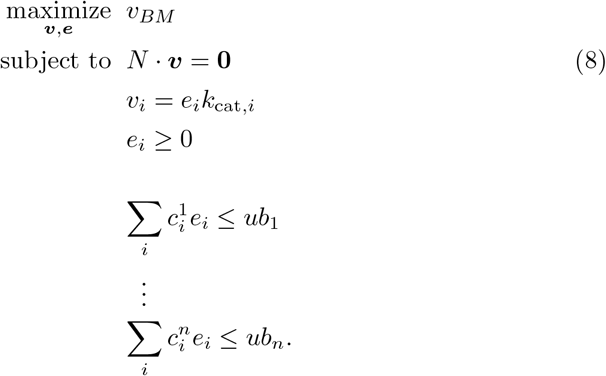

With the help of Equation (6) this problem can be solved with the reaction rates or with the enzyme concentrations as the optimization variables, as we show in SI3.3.

Basan et al. [27] made a core model that shows overflow metabolism in *E. coli* by dividing the proteome into three fractions: *φ*_*f*_, *φ*_*r*_ and *φ*_*BM*_, that thus sum up to one. These denote the fractions of the proteome catalyzing a fermentation, respiration and cell synthesis reaction, respectively, according to the relations: *v*_*f*_ = *ϵ*_*f*_ *φ*_*f*_, *v*_*r*_ = *ϵ_r_φ_r_*, and 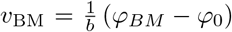. We will not define all unknown symbols in these and the following relations, since their interpretation is not relevant for this review. Note that the relation between the biomass reaction and the associated proteome fraction is non-standard, to include a non-growth associated maintenance term. Further, reactions for the uptake of a carbon source and the excretion of acetate^2^ are included, but these do not have an associated proteome fraction.

This gives the following steady state assumption

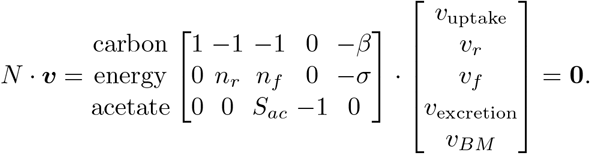

In addition, the authors set the uptake rate of nutrient: *v*_uptake_ = *c*_uptake_. Together, this yields a set of equations with only one solution; the variables (***v*** and ***ϕ***) can be directly calculated, and no optimization is required. The uniqueness of the solution is due to the small size of the model and because the constraints on uptake rate and the total proteome are modeled as equalities instead of inequalities. We show in SI3.4 that, for the appropriate choice of ***w***, *A*, ***x***, ***b***, this is equivalent to our standard form (1) in which the biomass production rate is maximized and the constraints are treated as inequalities:

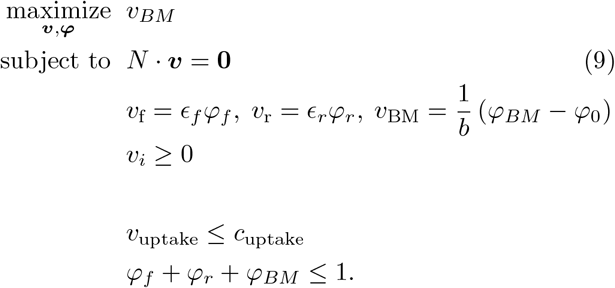

The authors assume that the yield of energy per carbon molecule is higher for respiration than for fermentation: *n*_*r*_ > *n*_*f*_^3^, but that fermentation is more proteome-efficient: *ϵ*_*f*_ > *ϵ*_*r*_. The enzyme cost of a certain reaction is the protein fraction necessary to attain one unit flux. We thus see that the enzyme costs of respiration are higher than fermentation: 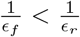. Because of this trade-off between yield and enzyme costs, it becomes optimal from a certain critical rate of carbon uptake to use the respiration and fermentation reactions simultaneously, so that the model shows overflow metabolism.

Vazquez et al. [24] responded to the explanation of Basan et al. by adding to the model that there is a maximum to the macromolecular density of a cell. They argue that the enzyme costs, as defined in the previous paragraph, should be proportional to the enzyme mass divided by its catalytic rate. The model that is used to explain overflow metabolism is thus the same, but with a different mechanistic underpinning of the *ϵ*-parameters. This reasoning was implemented earlier by the same authors in a genome-scale formalism called FBA with Macromolecular Crowding (FBAwMC), with which they already explained overflow metabolism in *E. coli* [23]. This formalism was later also used to model *S. cerevisiae* [29].

Another hypothesis is offered by Zhuang et al. [28] in which overflow metabolism is explained using a membrane occupancy constraint. The authors introduce parameters, *m*_*i*_, that capture the membrane area that is occupied per mol/liter of enzyme *i*. Assuming that there is only a limited plasma membrane budget, *B*_cyt_, this introduces the constraint ∑_*i*_*m*_*i*_*e*_*i*_ ≤ *B*_cyt_. Together with the steady state assumption, and a limited glucose uptake rate, this gives:

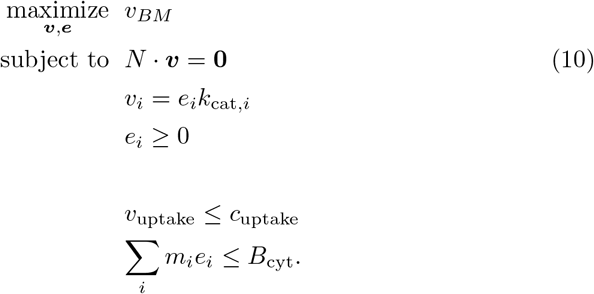

This hypothesis is supported by some quantitative evidence collected by Szenk et al. [22].

Note that the above resource allocation approaches differ in the mechanistic nature of the last constraint that is added, but that the optimization function, the steady state assumption and the limited substrate uptake are all similar. Shlomi et al. used a similar approach with a total proteome constraint to describe the Warburg effect in cancer cells [31].

A modeling approach that should be set slightly apart is Constrained Allocation FBA (CAFBA) [30]. The authors use only a total proteome constraint, and no direct limit on substrate uptake. Instead, substrate limitation is modeled by increasing a parameter *w*_*C*_ that captures the protein fraction needed for carbon catabolism to sustain one unit of carbon influx: *ϕ*_*C*_ = *ϕ*_*C*,0_ + *w*_C_*v*_*C*_. If the concentration of external nutrient decreases, *w*_*C*_ increases, and a larger fraction of the proteome is thus needed in the carbon catabolic sector. Because the sum of the proteome fractions needs to be one, this reduces the available proteome fraction for other sectors. As such, a change in nutrient concentration leads to a re-allocation of the proteome. The genome-scale model of *E. coli* can reproduce a switch from pure respiration to acetate secretion, but it does so with small discrete jumps. The gradual switch that is usually associated with overflow metabolism can only be found when the results are averaged over many different models created by choosing random parameters.

### 3.4 Self-fabrication models

In the previously described modeling approaches, the demand for cell components was approximated using a virtual biomass reaction. However, this approximation ignores an important nonlinear aspect of self-fabrication. A self-fabricating cell should produce two daughters identical to itself. The proportions in which the cell should produce cellular components thus depend on its own interior. If the cell reallocates resources to meet this demand for cellular components, its interior changes and therefore also the demand. The allocation of resources thus both depends on, and determines, the demand reaction.

Another inherent nonlinearity of cellular growth arises because cellular components dilute by growth: if a compound is not produced while the volume grows, its concentration drops. This dilution rate is equal to the growth rate of the cellular volume, so in steady state the net synthesis rate of all molecules should be equal to the growth rate. In turn, the same synthesis rates of all molecules determine how much volume is produced per unit time, and thus how fast the cell grows. The synthesis rate thus both depends on, and determines, the growth rate.

A small number of modeling approaches incorporate these two nonlinearities [7, 32–37]. The demand for cell synthesis components is calculated by the models instead of imposed on the models, and the growth rate can only be found after solving a nonlinear problem – or by solving a large number of linear problems in which the growth rate is treated as a parameter, as we will see. To keep our treatment of these complex models as accessible as possible, we will first describe the essential ingredients only. Then we will, referring to SI6 for most of the mathematical derivations, derive a set of relations that enables us to compare these self-fabrication models to the previously described models. After that, we will shortly discuss the various extensions that describe overflow metabolism.

The cell is modeled as consisting of three types of compounds: metabolites (with concentrations ***x***^4^ and possibly including macromolecules such as lipids or polynucleotides), enzymes (with concentrations ***e***), and the ribosome (with concentration *r*). The enzymes catalyze the conversion of metabolites into other metabolites. The ribosomes catalyze the synthesis of enzymes and ribosomes from metabolites. As before, it is assumed that the rates of the conversions scale proportionally with the concentrations of the catalysts, and kinetic saturation functions are again assumed constant:

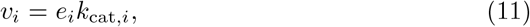

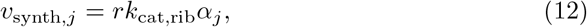

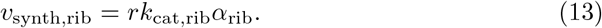

Here *v*_*i*_ is a usual metabolic reaction rate, and *v*_synth, *j*_ denotes the synthesis rate of enzyme *j*. The factor *α*_*j*_ is the fraction of the ribosome that is allocated to the synthesis of enzyme *j*, and since these are fractions we must have ∑_*j*_*α*_*j*_ = 1. It is further assumed that the concentrations of macromolecules add up to a fixed density^5^:

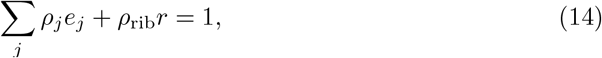

where the *ρ*_*j*_ are volumetric parameters.^6^ In a cell that is growing exponentially with rate *μ*, concentrations dilute with this same rate, see SI5 for a derivation. For the metabolites, this changes the steady state assumption from *N* · ***v*** = 0 in FBA approaches to *N* · ***v*** = *μ****x***. Moreover, if we explicitly model enzyme synthesis, we should also account for the metabolites that are consumed during this synthesis. Let *M* be the matrix that denotes how many metabolites are needed to make a specific enzyme, then we get a first set of constraints on the fluxs:

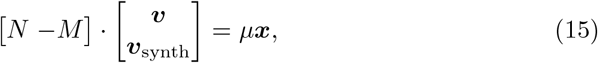

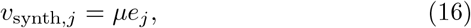

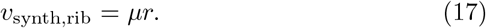

Equations (11) to (17) define the core ingredients of the self-fabrication models. In Supporting Information 6 we show how this system can be rewritten. In short: equations (11), (12) and (13) are used to get expressions for the concentrations ***e*** and *r* in terms of the fluxes, and these expressions are used in equations (14), (16) and (17) to get four relations between the fluxes ***v***, ***v***_synth_ and *μ*. These relations are linear in the reaction rates, so that the system can be written in a form that looks like a familiar Linear Program:

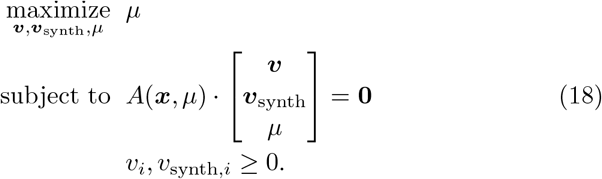

Although this looks like a Linear Program, it is more difficult since the constraint matrix is not constant: it depends on the metabolite concentrations, ***x***, and on the growth rate, *μ*. The ***x***-dependence is often “solved” by ignoring the dilution of small metabolites and fixing the concentrations of macromolecules based on experimental data [33,36]^7^. The *μ*-dependence that makes the problem nonlinear is overcome by fixing the growth rate in the constraint matrix to a certain value: *A*(***x***, *μ*) → *A*(***x***, *μ*_0_), and then add the constraint that the *μ* in the optimization variables should equal *μ*_0_. Note that, since it is fixed, we can no longer maximize the growth rate. However, we can check if there is a solution that solves the system. If there is no solution, then *μ*_0_ > *μ*_max_; if there is a solution, we can increase *μ*_0_. The maximal growth solution is found by repeating this procedure until the problem is still feasible for *μ*_0_ = *μ*_max_, but infeasible for all *μ*_0_ > *μ*_max_. So we get

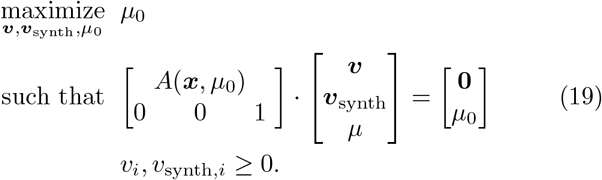

Using the described mathematical core, Goelzer et al. proposed a formalism that was named Resource Balance Analysis (RBA), with which they modeled overflow metabolism in *Bacillus subtilis* [38]. In addition to the density constraint of Equation (14), the authors used a constraint on the maximal concentration of macromolecules in the membrane; in our notation:∑_*j*_σ_*j*_*e*_*j*_ ≤ *D*_mem_.

In parallel, Thiele et al. [35] for *E. coli*, and Lerman et al. [34] for *Thermotoga maritima* presented the so-called Metabolism and Expression (ME) models. The mathematical basis of ME-models is equal to the basis of RBA-models (Equations (11) to (17)), but ME-models are even more comprehensive: for example, the synthesis rates of mRNA, tRNA, and RNA-polymerases are explicitly modeled. Moreover, some catalytic rates, of the ribosome for example, are no longer assumed to be independent of the growth rate; their dependence is estimated from experimental data. These extensions add many variables and constraints to the model, but we show in SI8 that these additions can still be written as relations that are linear in the reaction rates and nonlinear in the growth rate. In short, although the *A*-matrix of Equation (19) gets larger, ME-models can still be written in this form. O’Brien et al. used an ME-model to model overflow metabolism in *E. coli* [36]. The cytosolic density constraint was here supplemented with an upper bound on the substrate uptake flux.

Molenaar et al. [32] were the first to present a mechanistic model of cellular self-fabrication, a core model with 5 reactions and 3 metabolites. Because their model is so small, they could use enzyme kinetics and nonlinear optimization to directly maximize the growth rate. The optimal solutions show a discrete switch from an efficient pathway to an inefficient pathway. This is different from the gradual switch that is observed in overflow metabolism, even though, just as in [33], the authors model an upper bound on the membrane density. However, this does not effectively constrain the concentrations of membrane proteins, since the surface-to-volume ratio can be freely adjusted in the model. Therefore, only the density constraint of Equation (14) is effective. In personal correspondence, the authors confirmed that, in hindsight, it might have been more realistic to set a lower bound to the size of the cell. In this case, an additional constraint would have become active.

## 4 Discussion

### 4.1 Commonalities and differences

We reviewed many constrained optimization approaches that describe overflow metabolism, ranging from relatively simple linear Flux Balance Analysis models to complicated nonlinear Metabolism and Expression models. Some approaches use small core models, others use genome-scale networks comprising thousands of reactions. The imposed constraints are either limits on reaction rates, Gibbs dissipation limits or limits on enzyme concentrations. Despite all these differences, we managed to write all these models in a concise standard form by focusing on their mathematical essence. We conclude from this that these models must share a feature, one that must be essential for describing overflow metabolism. In the following, we will compare the models in their standard form using our recent theoretical work [7, 8] to analyze and explain this feature, using an extremum principle that governs the solutions of all the reviewed approaches.

#### 4.1.1 A general extremum principle: Overflow metabolism is caused by two growth-limiting constraints

All reviewed approaches model a growing cell by imposing a set of homogeneous constraints: a first set that ensures a steady state, and a second set that determines the feasible direction for irreversible reactions. There are some differences in how the first set is imposed. FBA and resource allocation approaches model a system that produces cell components in the proportions captured by a constant demand reaction, the biomass reaction. The steady state assumption ensures that no intermediate metabolite accumulates. The self-fabrication models implement this assumption with a balanced growth assumption: all cellular compounds should be produced to match the rate of consumption and dilution by growth. The demand reaction is therefore dependent on the growth rate, which gives rise to nonlinear relations between the optimization variables and the growth rate. These differences are illustrated in Figure 2.

**Fig. 2.**
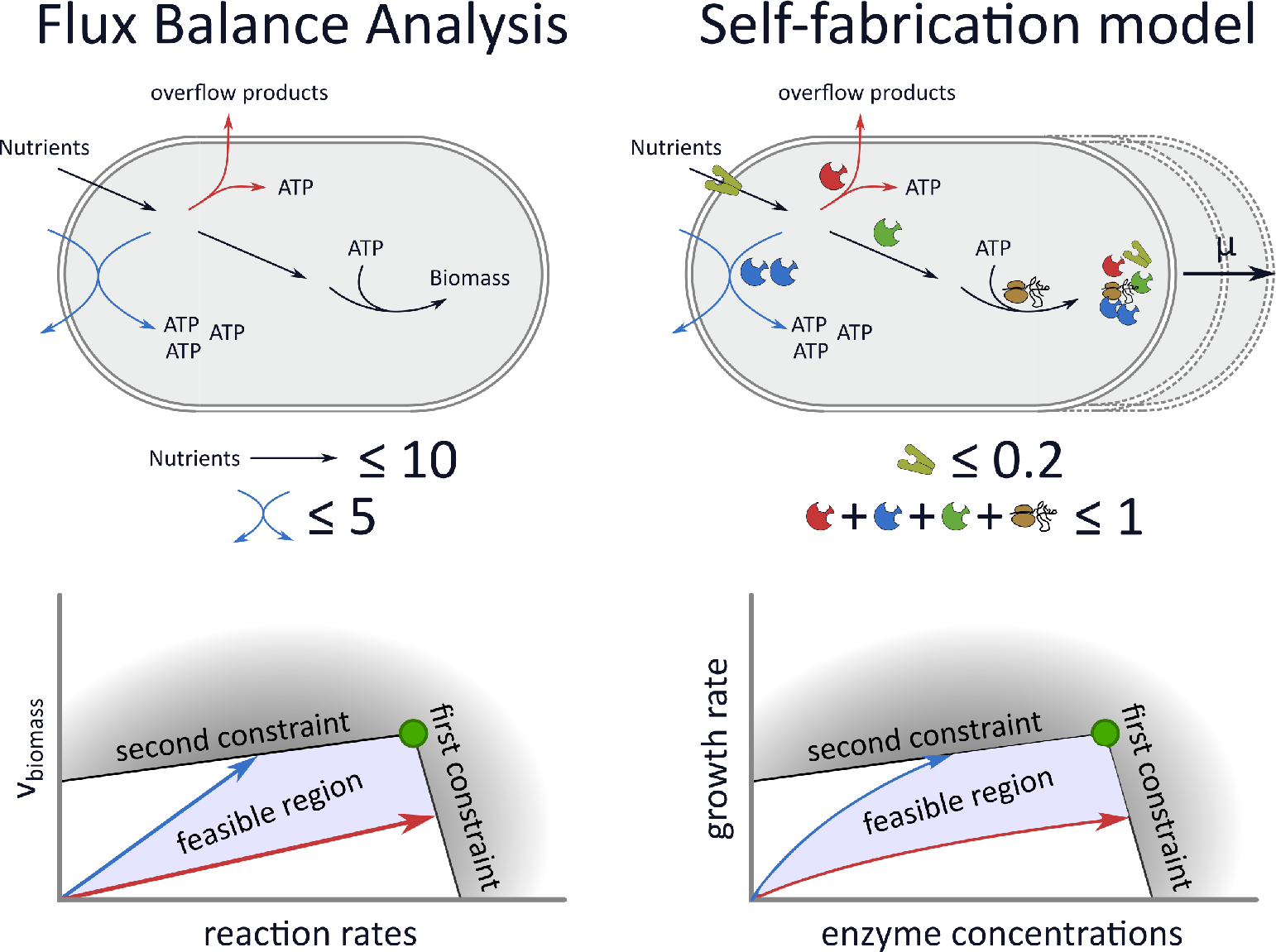
FBA models and self-fabrication models lead to a similar mathematical problem. In the **top** figures we illustrate two of the reviewed approaches. FBA models consider steady state fluxes through networks of metabolic reactions with constraints on the reaction rates. A virtual biomass reaction is added as a proxy for the growth rate. Self-fabricator models make the synthesis of enzymes and the ribosome explicit, and can therefore model the growth rate as the volume increase due to the production of components. The enzyme concentrations can now be viewed as the optimization variables, so that protein concentration constraints can also be included. In the **bottom** figures we show a highly simplified illustration of the solution space of both approaches. In the linear approaches, FBA and proteome-constrained models, all quantities depend linearly on the growth rate, while there are nonlinear dependencies in the self-fabricator models. However, we showed that in both cases, overflow metabolism is caused by two growth-limiting constraints.

Despite these seemingly different setups, we can define Elementary Modes in both cases: growth-supporting subnetworks that form the minimal building blocks of the solution space. These are called Elementary Flux Modes in the linear models [6], and Elementary Growth Modes in the self-fabricator models [7], see SI1 for a short introduction. The defining property of these modes is that all possible solutions of the growth models can be written as a combination of these modes. In other words, EFMs are non-decomposable metabolic subnetworks, and EGMs are non-decomposable self-fabrication subnetworks. Overflow metabolism is decomposable in an energy-efficient subnetwork, and a less energy-efficient subnetwork, and is thus a combination of two Elementary Modes.

Using the concept of Elementary Modes we derived an extremum principle stating that the number of flux-carrying Elementary Modes in the optimal solution will be smaller or equal than the number of active (i.e. growth-limiting) constraints. These growth-limiting constraints are the additional constraints that are imposed after the steady state and irreversibility constraints (this is illustrated and explained in Figure 1). The extremum principle implies that only one Elementary Mode will be selected by growth rate maximization under one constraint. For example, in a model in which only one nutrient uptake rate is constrained, we will never observe a gradual switch from a high-yield metabolic mode to the combination with a low-yield mode. Overflow metabolism must thus be a result of two constraints, see Figure 2 for an illustration of this result.

The extremum principle suggests that the success of describing overflow metabolism might not lie in the details of the stoichiometric networks, or the exact choices of model parameters, but rather in the mere existence of two constraints. This implies that finding the mechanistic cause of overflow metabolism amounts to finding which two constraints are actually limiting growth.

Unfortunately, the extremum principle does not predict which constraint causes overflow metabolism. It states that there should be two constraints, but does not reveal their identity. Moreover, overflow metabolism in different species might be due to completely different constraints. Thus, to find out which constraints cause overflow metabolism, we must test hypothetical constraints.

#### 4.1.2 Specific experiments: the mechanistic cause of overflow metabolism can be found with falsification experiments

Encouraged by the conclusion that the growth-limiting constraints must be important in causing overflow metabolism, we have listed all the constraints that are used in the reviewed models (Table 1). We see that almost all models indeed use two constraints.^8^ Molenaar et al. [32] and Mori et al. [30] form exceptions to this rule, using only one effective constraint. Our theory thus implies that only one EFM will be used in the optimal solutions of these models. Indeed, the models show discrete switches between EFMs when the growth rate increases. The model from Molenaar et al., containing only two EFMs, switches at once from respiration to fermentation. The genome-scale model of Mori et al. contains many different EFMs that form intermediate steps between full respiration and full fermentation. Their model therefore shows many small discrete switches, approximating a gradual switch.^9^ This gives rise to a separate hypothesis that we cannot fully exclude. However, we find it more probable that, upon a change in the environment, gene expression continuously tunes the proportions in which two EFMs are used, than that it repeatedly shuts down one EFM to upregulate another.

**Table 1:** An overview of the models that try to explain overflow metabolism, including which constraints were used in addition to the steady-state assumptions.

| Paper | Type | Constraint 1 | Constraint 2 |
| --- | --- | --- | --- |
| Majewski et al. [11] | FBA | glucose uptake rate | electron transfer capacity |
| Varma et al. [12] | FBA | glucose uptake rate | oxygen uptake rate |
| Carlson et al. [14] | FBA | glucose uptake rate | oxygen uptake rate |
| Niebel et al. [18] | tFBA | glucose uptake rate | free energy dissipation |
| Basan et al. [27] | resource | glucose uptake rate | total proteome |
| Mori et al. [30] | resource |  | total proteome |
| Vazquez et al. [23, 24] | resource | glucose uptake rate | macromolecular density |
| Van Hoek et al. [29] | resource | glucose uptake rate | macromolecular density |
| Zhuang et al. [28] | resource | glucose uptake rate | membrane occupancy |
| Szenk et al. [22] | resource | glucose uptake rate | membrane occupancy |
| Shlomi et al. [31] | resource | glucose uptake rate | total proteome |
| Molenaar et al. [32] | self-fabr |  | macromolecular density |
| Goelzer et al. [38] | self-fabr | membrane density | macromolecular density |
| O'Brien et al. [36] | self-fabr | glucose uptake rate | macromolecular density |

Among the models that use two constraints, there is some variation in the biological underpinnings of these constraints. The question is how to find the relevant constraints. The genome-scale approach is to make an extensive model and try to quantitatively match the experimental data. One risk, however, is overfitting, because a large enough model could potentially fit any experimental data. Such an approach should therefore be backed up by independent measurements of assumed constraints. Still, it is hard to imagine how a model could distinguish the effects caused by a ‘total proteome constraint’ and a ‘macromolecular density constraint’. For this, we need perturbation experiments. For example, to artificially perturb the proteome allocation of *E. coli*, Basan et al. [27, 39] overexpressed the nonfunctional protein LacZ in one experiment and added translation inhibitors in another. We have derived a formalism in which such perturbation experiments can be analyzed [8]. Basan et al. could provide evidence for their proposed total proteome constraint. In our opinion, this makes their proposed constraint the best-established mechanistic cause of overflow metabolism in *E. coli* up to this point. However, as we now know, there should be a second growth-limiting constraint. Basan et al. used a limit on the uptake rate of nutrients, which cannot truly be considered as a mechanistic cause of overflow metabolism, because, as described by Molenaar et al. in 2009: “… using an artificial maximal capacity constraint on substrate uptake ignores the possibility of variable investments made in substrate transport systems.” The identity of the second constraint in *E. coli*, even though a constraint on transport of glucose is generally used and thus apparently accepted, remains to be established.

#### 4.1.3 Towards a complete model of cellular self-fabrication

We observe two directions of development towards a complete model of cellular self-fabrication in the models that we have reviewed. Along the first direction the optimization variables are moved closer to the actual regulatory space of the cell, and thereby closer to the origin of overflow metabolism. Along the second direction, more and more of the inherent nonlinearity of self-fabrication is incorporated in the models.

To explain the first direction of development, we recall that FBA models use fluxes as variables, which cannot be directly regulated by the cell. Instead, enzyme concentrations are regulated and these will, together with the metabolite concentrations, determine the fluxes. The resource allocation models switch perspective to enzyme concentrations as variables with the major advantage that constraints on enzyme concentrations can be formulated directly. These constraints can be related to physically observable quantities, such as the available membrane area or cytosolic volume, whereas flux constraints cannot. Flux constraints can only be determined ad hoc using experimental data, which limits their predictive power.^10^ One can move even further towards the regulatory space of the cell because the enzyme concentrations are in fact dependent on the enzyme synthesis rates, and these are regu lated by the allocation of the ribosomes over the different mRNAs. The reviewed self-fabricator models indeed use as variables the enzyme synthesis rates [36, 38], or the ribosome allocation fractions [32]. A final step towards the regulatory space of the cell could be to model the regulation of mRNA synthesis via gene expression directly, but we do not know of any models that have implemented this.

The second direction of development moves towards incorporating three nonlinearities that are related to self-fabrication. We already mentioned two of them: 1) the dependence of the biomass composition on the enzyme allocation, and 2) the dependence of the demanded enzyme synthesis rates on the growth rate. The incorporation of these two nonlinearities form the main improvement of self-fabrication models with respect to FBA type models. Here we want to raise attention for a third nonlinearity: the kinetic dependence of enzyme and ribosome activities on the metabolite concentrations. If a cell reallocates resources, metabolite concentrations change as well, causing changes in the saturation levels of enzymes. Including the metabolite concentrations in the model however, requires information about the enzyme kinetics of all the different enzymes in the cell. Moreover, it makes the optimization problem computationally infeasible because the problem is no longer guaranteed to have only one local optimum. Therefore, global optimization software has to be used and it is difficult to ascertain that the found solution is the actual optimum. For these reasons, enzyme kinetics can so far only be included in core models [32] and theoretical work [7, 8, 37]. The question is if there are constraints, rules and patterns in the changes in (optimal) metabolite levels that would allow us to approximate optimal solutions without global optimisation of the full kinetic model.

#### 4.1.4 Alternatives for growth rate maximization

We have focused on growth rate maximization models, but alternative explanations of overflow metabolism cannot be fully excluded. For example, it might be that not absolute fitness is maximized, but rather relative fitness compared to competitors. For example, cells could produce overflow products to intoxicate their neighbours, or cells could maximize their uptake rate to claim the largest share of the nutrient pool. These explanations have been reviewed elsewhere [40, 41].

It might even be that cells are not completely optimized for anything. For example, it was shown that the overexpression of transcriptional regulator ArcA could increase the growth rate of *E. coli* on glycolytic substrates [42]. This shows that metabolism was not optimal in the wild type strain within the studied environmental conditions.

The sub-optimality of a population of microorganisms might be due to the high regulatory costs that would be required to steer each individual cell to the optimum. De Martino et al. [43] calculated a possible probability distribution for the metabolic states of single cells by maximizing the entropy of this distribution while the average growth rate was fixed to the measured value. This approach leads to a model of single-cell behaviour in which the least additional assumptions were made: ‘the probability distribution is as general as possible’. Their predicted distribution captured the measured fluxes better than a Flux Balance Analysis approach. Subsequently, they quantified the amount of regulation that would be needed to get a higher average growth rates, showing that attaining a maximal average growth rate would bring infinite regulatory costs.

## 5 Conclusion

We reviewed 15 different models of overflow metabolism, ranging from Flux Balance Analyses, to nonlinear self-fabricator models such as Metabolism and Expression models. Despite the many differences between the models, we could rewrite the mathematical cores of each of them into a concise standard form. This standard form could be analyzed using an extremum principle, stating that the number of Elementary Modes at maximal growth is less or equal than the number of growth-limiting constraints. The extremum principle implies that overflow metabolism is caused by at least two growth-limiting constraints. We therefore listed all reviewed models with their proposed constraints. We hope that this list will serve as a source of hypotheses that can now be tested using falsification experiments.

## Footnotes

1 We use a different standard form than the standard form that is used in Linear Programming. We found that our form better serves our purposes, but in SI2 we show that the two forms are equivalent.

2 The acetate excretion reaction was not explicitly mentioned in [27], but must have been included. We have made it explicit to be able to write a consistent stoichiometry matrix.

3 The authors originally used *e*_*r*_, *e*_*f*_ to denote these stoichiometric fractions, but we have renamed them to avoid confusion with enzyme concentrations.

4 The usage of ***x*** here is not related to its usage in our standard form, Equation (1)

5 In some modeling methods this density is modeled as an upper bound [32, 33, 37]. In SI7 we explain the advantages and disadvantages of doing this.

6 Dependent on the biochemical interpretation of the *ρ*-parameters, some models include contributions of the metabolite concentrations *ρ*_*i*_*x*_*i*_ [7, 37].

7 One could also solve the problem for fixed metabolite concentrations [7, 37], and then scan over all possible sets of concentrations, but this becomes computationally infeasible in large quantitative models.

8 Only Varma and Palsson use more than two constraints. Their third constraint is a lower bound on an ATP-maintenance reaction, which we have left out of the table for clarity.

9 The authors also present a figure showing a gradual switch, but this is the average behaviour for many models with slightly different parameters. The discontinuities are then averaged out.

10 In fact, many of the resource allocation models still use a flux constraint for the nutrient uptake reaction, so that these are also not entirely predictive.

