## Supplementary text for "The common message of constraint-based optimization approaches: overflow metabolism is caused by two growth-limiting constraints"

**Cellular Molecular Life Sciences manuscript No.**
(will be inserted by the editor)

### Supporting Information: The common message of constraint-based optimization approaches: overflow metabolism is caused by two growth-limiting constraints

**Daan H. de Groot · Julia Lischke · Riccardo Muolo · Robert Planqué · Frank J. Bruggeman · Bas Teusink**

Received: date / Accepted: date

#### 1 A short introduction to Elementary Modes

A growing cell converts nutrients into cellular building blocks via its metabolic networks, composed of hundreds to thousands of reactions and metabolites. It is often assumed that the cell grows at a fixed rate while its metabolism is at steady state, which means that the concentrations of all intracellular metabolites are constant in time. A steady cellular state can be imagined as a metabolic network through which a constant flux of matter flows from substrates to products. Not all network reactions need to be active in such a steady state.

An Elementary Flux Mode (EFM) is a minimal subnetwork (a set of reactions and metabolites) of the cell's metabolic network that is capable of sustaining a steady state. Thus, if we would remove all metabolic reactions that are not in the EFM, then there could still be a steady-state flux though the EFM. The minimality

Daan H. de Groot
Systems Bioinformatics, AIMMS, Vrije Universiteit Amsterdam, NL-1081HZ


Julia Lischke
Systems Bioinformatics, AIMMS, Vrije Universiteit Amsterdam, NL-1081HZ

Riccardo Muolo
Systems Bioinformatics, AIMMS, Vrije Universiteit Amsterdam, NL-1081HZ

Robert Planque
Department of Mathematics, Vrije Universiteit Amsterdam, NL-1081HV

Frank J. Bruggeman
Systems Bioinformatics, AIMMS, Vrije Universiteit Amsterdam, NL-1081HZ

Bas Teusink
Systems Bioinformatics, AIMMS, Vrije Universiteit Amsterdam, NL-1081HZ


means that if we would delete any reaction from the EFM, the steady state would be lost: some metabolite would accumulate or deplete.

A genome-scale network, typically containing a few thousand reactions, has an enormous number of these alternative pathways. There are so many EFMs, that currently available computational tools can not yet enumerate them all [1], so that we only know that there must be far more than billions of EFMs. Several EFMs can be active simultaneously, giving rise to a non-minimal subnetwork. Conversely, it has been shown [2] that any subnetwork that carries a steady state flux can be decomposed into EFMs. Therefore, the EFMs can be seen as the minimal building blocks of metabolic networks [2].

Now, let us consider a self-fabrication model: in addition to the metabolic network we also take into account enzyme synthesis by the ribosome and of the ribosome itself from metabolites. Again, we can assume a steady state, but now this means that the concentrations of enzymes and the ribosome should also be constant in time. Because cells grow, enzymes and ribosomes are diluted, and this dilution should be balanced by production[1]. This extended version of the steady state assumption is called the balanced growth assumption.

Balanced growth has an inherent nonlinearity. We can illustrate this with a thought experiment. Let us start with a set of enzyme and ribosome concentrations. The dilution rate of a compound is proportional to its concentration, and the concentrations of enzymes and ribosomes thus determine their own dilution rate. Their production rate should be equal to this dilution rate, because the concentrations should remain constant according to the balanced growth assumption. In turn, the metabolic reaction rates should be tuned to supply enough building blocks to match the production rate. However, the metabolic reaction rates are tuned by changing the concentrations of enzymes, so that we are back at the beginning of our thought experiment. Because of this circular problem, it is hard to find analogs of EFMs in whole cell-models: minimal pathways allowing self-fabrication.

Recently, we published a theory that studies such self-fabricating systems [3]. We identified minimal self-fabricating pathways, which we called *Elementary Growth Modes* (EGMs). These Elementary Modes are in many ways more cumbersome than EFMs, but conceptually they are very similar: in order to sustain balanced growth, no reaction can be removed. Thereby, any balanced growth state can be written as a sum of EGMs, confirming that EGMs are the minimal building blocks of self-fabrication models. Moreover, it has been proven that EFMs are an approximation of EGMs [3].

#### 2 Our standard form is equivalent to the standard Linear Programming form

In this section we will look into some mathematical details on Linear Programming (LP) that have been skipped in the main text. Let us start by introducing some notations

[1] The metabolites also dilute by growth, but this is usually ignored since the rates of production and consumption of metabolites are several orders of magnitude larger than the dilution rate

- given two column vectors $\boldsymbol{w}$ and $\boldsymbol{x}$ of $r$ components, the *scalar product* is defined in the following way

$$\boldsymbol{w}^T \cdot \boldsymbol{x} = \sum_{i=1}^{r} w_i x_i$$

where the row vector $\boldsymbol{w}^T$ is the *transpose* of the column vector $\boldsymbol{w}$, $w_i$ and $x_i$ are the $i^{\text{th}}$ components of vectors $\boldsymbol{w}$ and $\boldsymbol{x}$;
- given an $m \times r$ matrix $A$ with elements $a_{ij}$ and a column vector $\boldsymbol{x}$ of $r$ components, the matrix product between the two is defined in the following way

$$A \cdot \boldsymbol{x} = \boldsymbol{b}$$

where $\boldsymbol{b}$ is a column vector whose $m$ components are

$$b_i = \sum_{j=1}^{r} a_{ij} x_j$$

Let us also recall the ingredients of a linear programming optimization problem, as enumerated in the main text, i.e., we have the optimization variables $\boldsymbol{x}$, an objective function and constraints. The form in which those constraints are expressed defines whether the problem is written in the LP-standard form or not.

In this review we have chosen a form for the constraints ($A \cdot \boldsymbol{x} \leq \boldsymbol{b}$) that in our opinion is more intuitive and suits our framework better compared to the standard form ($A \cdot \boldsymbol{x} = \boldsymbol{b}$), even though certain constraints found in literature are expressed in the latter form, e.g., the steady state constraint. In the following, we will show that these forms are mathematically equivalent.

Before proceeding: it is customary to write linear programming problems in the form of *minimizing* a certain objective function, while sometimes this function is *maximized*. As the maximizers of a certain function $\boldsymbol{w}^T \cdot \boldsymbol{x}$ are also the minimizers of $(-\boldsymbol{w}^T) \cdot \boldsymbol{x}$, the two approaches are equivalent. Because we generally assume that cells are forced to reproduce as fast as possible to outcompete the rest of the population, we have chosen the approach where the objective function is maximized.

**Theorem 1** *Let $\boldsymbol{w}$, $\boldsymbol{x}$ and $\boldsymbol{b}$ be column vectors of respectively $r$, $r$ and $m$ components and let $A$ be an $m \times r$ matrix. The linear programming problem posed in 'our standard form':*

$$\begin{aligned} \underset{\boldsymbol{x}}{\text{maximize}} \quad & \boldsymbol{w}^T \cdot \boldsymbol{x} \\ \text{subject to} \quad & A \cdot \boldsymbol{x} \leq \boldsymbol{b} \\ & x_i \geq 0 \end{aligned} \tag{1}$$

*is equivalent to a problem in the 'LP-standard form':*

$$\begin{aligned} \underset{\boldsymbol{x}}{\text{maximize}} \quad & \boldsymbol{y}^T \cdot \boldsymbol{\xi} \\ \text{subject to} \quad & \mathcal{A} \cdot \boldsymbol{\xi} = \boldsymbol{b} \\ & \xi_i \geq 0, \end{aligned} \tag{2}$$

*for suitable choices of $y$, $\xi$, and $\mathcal{A}$.*

*Proof* We start from the problem in our form

$$\begin{aligned} \underset{\boldsymbol{x}}{\text{maximize}} \quad & \boldsymbol{w}^T \cdot \boldsymbol{x} \\ \text{subject to} \quad & A \cdot \boldsymbol{x} \leq \boldsymbol{b} \\ & x_i \geq 0. \end{aligned}$$

Note that $A \cdot \boldsymbol{x} \leq \boldsymbol{b}$ can also be expressed as that there is room for a vector with positive entries that is the difference between $\boldsymbol{b}$ and $A \cdot \boldsymbol{x}$. We introduce this vector $\boldsymbol{s}$ of $m$ components that are the *slack variables*, defined as

$$\boldsymbol{s} = \boldsymbol{b} - A \cdot \boldsymbol{x},$$

so that

$$A \cdot \boldsymbol{x} + \boldsymbol{s} = \boldsymbol{b}.$$

The problem in Equation (1) becomes

$$\begin{aligned} \underset{\boldsymbol{x}}{\text{maximize}} \quad & \boldsymbol{w}^T \cdot \boldsymbol{x} \\ \text{subject to} \quad & A \cdot \boldsymbol{x} + \boldsymbol{s} = \boldsymbol{b} \\ & x_i \geq 0 \\ & s_i \geq 0. \end{aligned}$$

If we define new variables

$$\boldsymbol{y} = \begin{bmatrix} \boldsymbol{w} \\ \boldsymbol{0} \end{bmatrix} \quad , \quad \boldsymbol{\xi} = \begin{bmatrix} \boldsymbol{x} \\ \boldsymbol{s} \end{bmatrix} \quad , \quad \mathcal{A} = \left[ A \mid \mathbb{I}_{m \times m} \right],$$

where $\boldsymbol{y}$ and $\boldsymbol{\xi}$ are column vectors of $n + m$ components, $\mathcal{A}$ is an $m \times (n + m)$ matrix and $\mathbb{I}_{m \times m}$ is the $m \times m$ identity matrix, we can rewrite the above problem in standard form

$$\begin{aligned} \underset{\boldsymbol{\xi}}{\text{maximize}} \quad & \boldsymbol{y}^T \cdot \boldsymbol{\xi} \\ \text{subject to} \quad & \mathcal{A} \cdot \boldsymbol{\xi} = \boldsymbol{b} \\ & \xi_i \geq 0. \end{aligned}$$

If we find a solution for this problem, this can be directly mapped to a solution for the problem in Equation (1). So, we have proven that all problems written in our form can be written in the standard LP-form.

The converse can also be proven. Suppose we have a linear programming problem written in LP-standard form:

$$\begin{aligned} \underset{\boldsymbol{x}}{\text{maximize}} \quad & \boldsymbol{w}^T \cdot \boldsymbol{x} \\ \text{subject to} \quad & A \cdot \boldsymbol{x} = \boldsymbol{b} \\ & x_i \geq 0. \end{aligned}$$

Note that the constraint $A \cdot \boldsymbol{x} = \boldsymbol{b}$ is equivalent to demanding that $A \cdot \boldsymbol{x} \leq \boldsymbol{b}$ and $A \cdot \boldsymbol{x} \geq \boldsymbol{b}$. In turn, the latter is equivalent to the constraint: $-A \cdot \boldsymbol{x} \leq -\boldsymbol{b}$. If we now choose $\mathcal{A} = \begin{bmatrix} A \\ -A \end{bmatrix}$ and $\boldsymbol{\beta} = \begin{bmatrix} \boldsymbol{b} \\ -\boldsymbol{b} \end{bmatrix}$, the problem can be rewritten as

$$\begin{aligned} \underset{\boldsymbol{x}}{\text{maximize}} \quad & \boldsymbol{w}^T \cdot \boldsymbol{x} \\ \text{subject to} \quad & \mathcal{A} \cdot \boldsymbol{x} \leq \boldsymbol{\beta} \\ & x_i \geq 0 \end{aligned}$$

With this we have proven that all Linear Programs written in LP-standard form can be rewritten in our form, completing the proof.

#### 3 The standard form for the different types of models

##### 3.1 The standard form for FBA models

The general form for an FBA problem is the following

$$\begin{aligned} \underset{\boldsymbol{v}}{\text{maximize}} \quad & v_{BM} \\ \text{subject to} \quad & N \cdot \boldsymbol{v} = \mathbf{0} \\ & ub_i \geq v_i \geq lb_i \geq 0. \end{aligned}$$

We can rewrite the above FBA problem to fit our standard form of Equation (1), by, amongst others, using the equivalence of $v_i \geq lb_i$ to $-vi \leq -lb_i$. We define

$$\boldsymbol{x} = \begin{bmatrix} v_1 \\ \vdots \\ v_{r-1} \\ v_{BM} \end{bmatrix} \quad \boldsymbol{w} = \begin{bmatrix} 0 \\ \vdots \\ 0 \\ 1 \end{bmatrix} \quad \mathcal{N} = \begin{bmatrix} N \\ -N \\ \mathbb{I}_{r\times r} \\ -\mathbb{I}_{r\times r} \end{bmatrix}$$

$$\boldsymbol{b}_1 = \begin{bmatrix} 0 \\ \vdots \\ 0 \end{bmatrix} \quad \boldsymbol{b}_2 = \begin{bmatrix} 0 \\ \vdots \\ 0 \end{bmatrix} \quad \boldsymbol{b}_3 = \begin{bmatrix} ub_1 \\ \vdots \\ ub_r \end{bmatrix} \quad \boldsymbol{b}_4 = \begin{bmatrix} -lb_1 \\ \vdots \\ -lb_r \end{bmatrix}$$

and lastly,

$$\boldsymbol{\beta} = \begin{bmatrix} \boldsymbol{b}_1 \\ \boldsymbol{b}_2 \\ \boldsymbol{b}_3 \\ \boldsymbol{b}_4 \end{bmatrix}$$

we obtain our standard form

$$\begin{aligned} \underset{\boldsymbol{x}}{\text{maximize}} \quad & \boldsymbol{w}^T \cdot \boldsymbol{x} \\ \text{subject to} \quad & \mathcal{N} \cdot \boldsymbol{x} \leq \boldsymbol{\beta} \\ & x_i \geq 0. \end{aligned}$$

3.2 The standard form for the model of Niebel et al.

In the main text, we reduced the thermodynamic-FBA model of Niebel et al. [4] to the following form

$$\begin{aligned}
\underset{\boldsymbol{v}}{\text{maximize}} \quad & v_{BM} \\
\text{subject to} \quad & N \cdot \boldsymbol{v} = \boldsymbol{0} \\
& v_j \geq 0 \\
& \Delta_r G'_j(\boldsymbol{c}) < 0 \quad \text{for all reactions with } v_j \neq 0 \\
\\
& v_{\text{Glc,uptake}} \leq b_{\text{Glc}} \\
& -\sum_j \Delta_r G'_j(\boldsymbol{c}) v_j \leq g^{\text{diss}}_{\text{lim}}.
\end{aligned} \tag{3}$$

We start by looking at the constraint: $\Delta_r G'_j(\boldsymbol{c}) < 0$ for all reactions with $v_j \neq 0$. Since linear programs cannot work with strict inequalities, the authors approximated this constraint by: $\Delta_r G'_j(\boldsymbol{c}) \leq -0.5$. This inequality should however only be restrictive if $v_j$ is nonzero. We can incorporate this by multiplying both sides by $v_j$: if $v_j = 0$ then the inequality holds trivially, while the constraint becomes restrictive if $v_j \neq 0$. Thus we can write $\Delta_r G'_j(\boldsymbol{c}) v_j \leq -0.5 v_j$.

The other constraints can be incorporated more easily. We define

$$\boldsymbol{x} = \begin{bmatrix} v_{\text{Glc,uptake}} \\ \vdots \\ v_{r-1} \\ v_{BM} \end{bmatrix} \qquad \boldsymbol{w} = \begin{bmatrix} 0 \\ \vdots \\ 0 \\ 1 \end{bmatrix} \qquad \mathcal{N} = \begin{bmatrix} N \\ -N \\ -\mathcal{A} \\ \mathcal{B} \end{bmatrix}$$

where

$$\mathcal{A} = \begin{bmatrix} \Delta_r G'_1(\boldsymbol{c}) + 0.5 & \cdots & 0 \\ \vdots & \ddots & \vdots \\ 0 & \cdots & \Delta_r G'_r(\boldsymbol{c}) + 0.5 \end{bmatrix} \qquad \mathcal{B} = \begin{bmatrix} -\Delta_r G'_1(\boldsymbol{c}) & \cdots & -\Delta_r G'_r(\boldsymbol{c}) \end{bmatrix}$$

and

$$\boldsymbol{b}_1 = \begin{bmatrix} 0 \\ \vdots \\ 0 \end{bmatrix} \quad \boldsymbol{b}_2 = \begin{bmatrix} 0 \\ \vdots \\ 0 \end{bmatrix} \quad \boldsymbol{b}_3 = \begin{bmatrix} 0 \\ \vdots \\ 0 \end{bmatrix} \quad \boldsymbol{b}_4 = g^{\text{diss}}_{\text{lim}}$$

Then, by defining

$$\boldsymbol{\beta} = \begin{bmatrix} \boldsymbol{b}_1 \\ \boldsymbol{b}_2 \\ \boldsymbol{b}_3 \\ \boldsymbol{b}_4 \end{bmatrix}$$

we obtain our standard form

$$\begin{aligned}
\underset{\boldsymbol{x}}{\text{maximize}} \quad & \boldsymbol{w}^T \cdot \boldsymbol{x} \\
\text{subject to} \quad & \mathcal{N} \cdot \boldsymbol{x} \leq \boldsymbol{\beta} \\
& x_i \geq 0.
\end{aligned}$$

##### 3.3 The standard form for resource allocation models

The general form for the resource allocation modeling approach is the following

$$\begin{aligned} \underset{\boldsymbol{v},\boldsymbol{e}}{\text{maximize}} \quad & v_{BM} \\ \text{subject to} \quad & N \cdot \boldsymbol{v} = \mathbf{0} \\ & v_i = e_i k_{\text{cat},i} \\ & e_i \geq 0 \\ & \sum_i c_i^1 e_i \leq ub_1 \\ & \vdots \\ & \sum_i c_i^n e_i \leq ub_n. \end{aligned} \tag{4}$$

This model is a resource allocation model, so essentially the enzyme concentrations should be the optimization variables, since these are the resources that are allocated. In the main text we however chose to express the steady state condition in terms of the fluxes, since this is customary in literature. Fortunately, we can rewrite the problem using enzyme concentrations as optimization variables, by using the relation between enzyme concentrations and fluxes:$v_i = e_i k_{\text{cat},i}$. We define a new stoichiometry matrix $\hat{N}$ as follows:

$$\hat{N} = \left[k_{\text{cat},1} N_1 \, \ldots \, k_{\text{cat},r} N_r\right],$$

where $N_i$ are the columns of the stoichiometry matrix $N$. Furthermore, the rate of the biomass reaction is written as $v_{BM} = e_{BM} k_{\text{cat},BM}$.[2] Then, we can rewrite the linear programming problem in an equivalent form, where now the optimization varibles are the enzymes

$$\begin{aligned} \underset{\boldsymbol{e}}{\text{maximize}} \quad & e_{BM} k_{\text{cat},BM} \\ \text{subject to} \quad & \hat{N} \cdot \boldsymbol{e} = \mathbf{0} \\ & \sum_i c_i^1 e_i \leq ub_1 \\ & \vdots \\ & \sum_i c_i^n e_i \leq ub_n. \end{aligned}$$

[2] It depends on the modeling method if the enzyme $e_{BM}$ is considered real or virtual. In case it is considered virtual, we can still model it in this way but we should add a constraint that forces it to be 1 and choose $k_{\text{cat},BM} = 1$

This gives us a formulation with the objective function and all constraints expressed in the same variables. We define

$$\boldsymbol{x} = \begin{bmatrix} e_1 \\ \vdots \\ e_{r-1} \\ e_{BM} \end{bmatrix} \qquad \boldsymbol{w} = \begin{bmatrix} 0 \\ \vdots \\ 0 \\ k_{\text{cat},BM} \end{bmatrix} \qquad \mathcal{N} = \begin{bmatrix} N \\ -N \\ \mathcal{A} \end{bmatrix},$$

where

$$\mathcal{A} = \begin{bmatrix} c_1^1 & \cdots & c_r^1 \\ \vdots & \ddots & \vdots \\ c_1^n & \cdots & c_r^n \end{bmatrix},$$

and

$$\boldsymbol{b}_1 = \begin{bmatrix} 0 \\ \vdots \\ 0 \end{bmatrix} \qquad \boldsymbol{b}_2 = \begin{bmatrix} 0 \\ \vdots \\ 0 \end{bmatrix} \qquad \boldsymbol{b}_3 = \begin{bmatrix} ub_1 \\ \vdots \\ ub_r \end{bmatrix} \quad \text{and} \quad \boldsymbol{\beta} = \begin{bmatrix} \boldsymbol{b}_1 \\ \boldsymbol{b}_2 \\ \boldsymbol{b}_3 \end{bmatrix}.$$

We obtain our standard form

$$\begin{aligned} \underset{\boldsymbol{x}}{\text{maximize}} \quad & \boldsymbol{w}^T \cdot \boldsymbol{x} \\ \text{subject to} \quad & \mathcal{N} \cdot \boldsymbol{x} \leq \boldsymbol{\beta} \\ & x_i \geq 0. \end{aligned}$$

3.4 The standard form for the model of Basan et al. [5]

In the main text we derived that Basan et al. [5] solve the following problem:

$$\begin{aligned} & N \cdot \boldsymbol{v} = \boldsymbol{0} \\ & v_i \geq 0 \\ & v_{\text{uptake}} = c_{\text{uptake}} \\ & \varphi_f + \varphi_r + \varphi_{BM} = 1, \end{aligned} \tag{5}$$

where

$$N = \begin{bmatrix} 1 & -1 & -1 & 0 & -\beta \\ 0 & n_r & n_f & 0 & -\sigma \\ 0 & 0 & S_{ac} & -1 & 0 \end{bmatrix} \quad , \quad \boldsymbol{v} = \begin{bmatrix} v_{\text{uptake}} \\ v_r \\ v_f \\ v_{excretion} \\ v_{BM} \end{bmatrix}$$

and

$$v_{\text{f}} = \epsilon_f \varphi_f \ , \quad v_{\text{r}} = \epsilon_r \varphi_r \ , \quad v_{\text{BM}} = \frac{1}{b} \left( \varphi_{BM} - \varphi_0 \right). \tag{6}$$

This gives a set of five equalities: three equalities are due to assuming a steady-state, one is due to setting the uptake rate, and one comes from assuming that the proteome fractions add up to 1. These five equalities completely determine the five reaction rates. There is therefore only one solution, which is thus automatically the optimal solution and no further optimization is required. However, this model can be rewritten as a constraint-based optimization if we view the constaints on uptake and on the proteome fractions as inequalities. This will not affect the solutions, since the biomass production rate will only be maximized if these constraints are maximally exploited: a higher uptake rate and more protein investment will always lead to a higher production rate. We can thus write

$$\begin{aligned} \underset{\boldsymbol{v}}{\text{maximize}} \quad & v_{BM} \\ \text{subject to} \quad & N \cdot \boldsymbol{v} = \boldsymbol{0} \\ & v_i \geq 0 \\ & v_{\text{uptake}} \leq c_{\text{uptake}} \\ & \varphi_f + \varphi_r + \varphi_{BM} \leq 1. \end{aligned} \tag{7}$$

This model could also be written with the proteome fractions as the optimization variables, but that would involve introducing proteome fractions corresponding to the uptake and excretion reactions, which were ignored by the authors. We will therefore keep using fluxes as the variables. The constraint $\varphi_r + \varphi_f + \varphi_{BM} \leq 1$ is rewritten using the relations in Equation (6). We get

$$\frac{1}{\epsilon_r} v_r + \frac{1}{\epsilon_f} v_f + b v_{BM} + \phi_0 \leq 1.$$

This can be written in our standard form by defining

$$\boldsymbol{x} = \begin{bmatrix} v_{\text{uptake}} \\ v_r \\ v_f \\ v_{BM} \\ v_{excretion} \end{bmatrix} \qquad \boldsymbol{w} = \begin{bmatrix} 0 \\ 0 \\ 0 \\ 0 \\ 1 \end{bmatrix} \qquad \mathcal{N} = \begin{bmatrix} N \\ -N \\ \mathcal{A} \end{bmatrix}$$

where

$$\mathcal{A} = \begin{bmatrix} 1 & 0 & 0 & 0 & 0 \\ 0 & \frac{1}{\epsilon_r} & \frac{1}{\epsilon_f} & b & 0 \\ 0 & 0 & 0 & 0 & 0 \\ 0 & 0 & 0 & 0 & 0 \end{bmatrix}$$

and

$$\boldsymbol{b}_1 = \boldsymbol{b}_2 = \begin{bmatrix} 0 \\ 0 \\ 0 \end{bmatrix} \qquad \boldsymbol{b}_3 = \begin{bmatrix} c_{\text{uptake}} \\ 1 - \varphi_0 \\ 0 \\ 0 \end{bmatrix} \quad \text{and} \quad \boldsymbol{\beta} = \begin{bmatrix} \boldsymbol{b}_1 \\ \boldsymbol{b}_2 \\ \boldsymbol{b}_3 \end{bmatrix}$$

By means of the newly defined matrices and vectors, we get

$$\begin{aligned} \underset{\boldsymbol{x}}{\text{maximize}} \quad & \boldsymbol{w}^T \cdot \boldsymbol{x} \\ \text{subject to} \quad & \mathcal{N} \cdot \boldsymbol{x} \leq \boldsymbol{\beta} \\ & x_i \geq 0. \end{aligned}$$

#### 4 Mass conservation should imply conservation of Gibbs energy

In the thermodynamic FBA approach by Niebel et al. [4] a constraint is imposed that we believe is equivalent to the steady state constraint. Therefore, we think that only one of the two constraints has to be imposed.

The constraint is a Gibbs energy balance:

$$g^{\text{diss}} = \sum_{i \in \text{EXG}} g_i = \sum_{j \in \text{MET}} g_j, \tag{8}$$

where MET is a set of all metabolic processes in the cell, EXG is the set of all exchange processes, and $g_i$ denotes the Gibbs energy exchange rate for $i \in$ EXG and the Gibbs energy dissipation rate for $j \in$ MET.

At this point we refer to a review by von Stockar et al. [6]. Equation (3) in this review states that for a cell with a steady state metabolism, the following holds:

$$\frac{dG}{dt} = \dot{W} + \sum_i \mu_i \dot{n}_i - \mu_{BM} \dot{n}_{BM} - T\dot{S}_{\text{prod}}. \tag{9}$$

Here, $\frac{dG}{dt}$, denotes the change in Gibbs free energy in the cell. Because the cell is assumed to be in steady state, this derivative should be zero. The first term on the right hand side, $\dot{W}$, is the external non-chemical work done on the system. The authors do not mention any possible sources of such work so that this term should also be zero. The second and third term, are the exchange rates of Gibbs free energy with the environment in the form of substrates/products and biomass, respectively. These two terms are equal to $\sum_{i \in \text{EXG}} g_i$ in the constraint imposed by Niebel et al. The fourth therm, $T\dot{S}_{\text{prod}}$ is equal to the Gibbs energy that is dissipated by the cell's internal processes. This is equal to the term $\sum_{j \in \text{MET}} g_j$ from Equation (8).

To satisfy Equation (9), we must have that the fourth term is equal to the sum of the second and third term, as long as no non-chemical work is done. This, however, would immediately imply that (8) holds. We thus see that, the constraint that is imposed by Niebel et al. is always satisfied by a cell in steady state, as long as no work is done.

We therefore believe that the Gibbs energy balance is always satisfied and thus redundant in the modeling method.

##### 4.1 A more detailed investigation

We can also start from the precise definitions of the various parameters in the constraint, as given by the authors in the Supporting Information. We will expand Equation (8) to see if all terms cancel if we assume a steady state. The Gibbs energy exchange rates are defined by

$$g_i = \Delta_f G'_i v_i \qquad \text{for } i \in \text{EXG}, \tag{10}$$

$$g_j = \Delta_r G'_j v_j \qquad \text{for } j \in \text{MET}, \tag{11}$$

where $v_i$ denotes a reaction rate, $\Delta_f G'_i$ the Gibbs energy of formation of the exchanged metabolites, and $\Delta_r G'_j$ the Gibbs energy of reaction $j$. These are in turn given by

$$\Delta_f G'_i = \Delta_f G'^0_i + RT\ln c_i \qquad \text{for } i \in \text{EXG}, \tag{12}$$
$$\Delta_r G'_j = \Delta_r G'^0_j + \Delta_r G'^t_j + RT\textstyle\sum_{i\notin h^+} S_{ij}\ln c_i \qquad \text{for } j \in \text{MET}. \tag{13}$$

Before defining the new variables introduced here, we can fill in the formula for $\Delta_r G'^0_j$, giving

$$\Delta_f G'_i = \Delta_f G'^0_i + RT\ln c_i \qquad \text{for } i \in \text{EXG}, \tag{14}$$
$$\Delta_r G'_j = \textstyle\sum_{i\notin h^+} S_{ij}\Delta_f G'^0_i + \Delta_r G'^t_j + RT\sum_{i\notin h^+} S_{ij}\ln c_i \qquad \text{for } j \in \text{MET} \tag{15}$$

where $\Delta_f G'^0_i$ is the standard Gibbs energy of formation of reactant $i$, and $\Delta_r G'^t_j$ is the Gibbs energy of metabolite transport. The latter is given by

$$\Delta_r G'^t_j = RT\sum_{\iota\notin h^+} s_{\iota j}\gamma_\iota + RT s_{h^+[in]j}\left(\ln c_{h^+[in]} - \ln c_{h^+[out]}\right) + F S_{Q[in]j}\Delta\phi_j. \tag{16}$$

In addition to these definitions, the authors use the following steady state constraint:

$$\sum_{j\in\text{MET}} S_{ij}v_j = v_{i\in\text{EXG}}, \tag{17}$$

where $v_{i\in\text{EXG}}$ denotes the exchange of metabolite $i$ over the system boundary. This relation is thus $\sum_{j\in\text{MET}} S_{ij}v_j = 0$ for $i$ an internal metabolite, since exchange over the system boundary cannot occur from within the cell.

We now put everything back together in Equation (8), although we do not expand the Gibbs energy of metabolite transport for now. The sum over all exchange processes gives

$$\begin{aligned}\sum_{i\in\text{EXG}} g_i &= \sum_{i\in\text{EXG}} \Delta_f G'_i v_i,\\ &= \sum_{i\in\text{EXG}}\left(\Delta_f G'^0_i + RT\ln c_i\right)v_i.\end{aligned}$$

The sum over all metabolic processes in the cell becomes

$$\begin{aligned}\sum_{j\in\text{MET}} g_j &= \sum_{j\in\text{MET}} \Delta_r G'_j v_j,\\ &= \sum_{j\in\text{MET}}\left(\sum_{i\notin h^+} S_{ij}\Delta_f G'^0_i + \Delta_r G'^t_j + RT\sum_{i\notin h^+} S_{ij}\ln c_i\right)v_j,\\ &= \sum_{j\in\text{MET}}\left(\sum_i S_{ij}\Delta_f G'^0_i + \Delta_r G'^t_j + RT\sum_i S_{ij}\ln c_i\right)v_j\\ &\quad - \sum_{j\in\text{MET}}\left(S_{h^+j}\Delta_f G'^0_{h^+} + RT S_{h^+j}\ln c_{h^+}\right)v_j.\end{aligned}$$

We can exchange the order of the sums and move all the terms that are not dependent on $j$ out of the sum over $j$. Subsequently using the steady state assumption in Equation (17) we get

$$\begin{aligned}\sum_{j\in\text{MET}} g_j &= \sum_i \Delta_f G_i'^0 \left(\sum_{j\in\text{MET}} S_{ij} v_j\right) + \sum_{j\in\text{MET}} \Delta_r G_j'^t v_j + RT \sum_i \ln c_i \left(\sum_{j\in\text{MET}} S_{ij} v_j\right) \\ &\quad - \sum_{j\in\text{MET}} \left(S_{h^+j} \Delta_f G_{h^+}'^0 + RT S_{h^+j} \ln c_{h^+}\right) v_j, \\ &= \sum_i \Delta_f G_i'^0 v_i + \sum_{j\in\text{MET}} \Delta_r G_j'^t v_j + RT \sum_i \ln c_i v_i \\ &\quad - \sum_{j\in\text{MET}} \left(S_{h^+j} \Delta_f G_{h^+}'^0 + RT S_{h^+j} \ln c_{h^+}\right) v_j, \\ &= \sum_i \left(\Delta_f G_i'^0 + RT \sum_i \ln c_i\right) v_i + \sum_{j\in\text{MET}} \Delta_r G_j'^t v_j \\ &\quad - \sum_{j\in\text{MET}} \left(S_{h^+j} \Delta_f G_{h^+}'^0 + RT S_{h^+j} \ln c_{h^+}\right) v_j.\end{aligned}$$

Comparing this to the sum of Gibbs energy exchange, we see that many terms will cancel. We get

$$g_{\text{EXG}}^{\text{diss}} - g_{\text{MET}}^{\text{diss}} = \sum_{j\in\text{MET}} \left(S_{h^+j} \Delta_f G_{h^+}'^0 + RT S_{h^+j} \ln c_{h^+}\right) v_j - \sum_{j\in\text{MET}} \Delta_r G_j'^t v_j = 0.$$

Or, written diffently,

$$\sum_{j\in\text{MET}} \left(S_{h^+j} \Delta_f G_{h^+}'^0 + RT S_{h^+j} \ln c_{h^+}\right) v_j = \sum_{j\in\text{MET}} \Delta_r G_j'^t v_j. \tag{18}$$

This can be expanded further by using Equation (16):

$$\begin{aligned}&\sum_{j\in\text{MET}} \left(S_{h^+j} \Delta_f G_{h^+}'^0 + RT S_{h^+j} \ln c_{h^+}\right) v_j = \\ &\qquad \sum_{j\in\text{MET}} \left(RT \sum_{\iota\notin h^+} s_{\iota j}\gamma_\iota + RT s_{h^+[in]j} \left(\ln c_{h^+[in]} - \ln c_{h^+[out]}\right) + F S_{Q[in]j} \Delta\phi_j\right) v_j.\end{aligned}$$

Niebel et al. state that the three contributions of the Gibbs energies changes of transport are due to "(i) the transport of species $\iota$ between compartments with different pH values and the concomitant release or binding of protons caused by the protonation or de-protonation of the transported species, (ii) the translocations of protons by proton sym-/antiporters or proton pumps; (iii) the transport of charged metabolites across electrical membrane potentials".

We do not see how this can be simplified any further, but if indeed we are wrong in believing that the Gibbs energy balance is equivalent to a mass balance, then the difference should reside in this last equality. Then, the Gibbs energy balance would give an equality between the Gibbs energy dissipated in reactions involving protons, and the Gibbs energy involved in transport processes. We do not entirely understand what this means, but we have contacted the authors to investigate this further.

#### 5 Cellular compounds dilute with rate $\mu$

We consider a cell of which the volume grows exponentially with rate $\mu$: $\mu = \frac{1}{V}\frac{dV}{dt}$. The cell contains a compound $X$ with copy number $n_x$ and concentration $x = \frac{n_x}{V}$. We assume that this compound is not synthesized, so that its copy number is constant in time: $\frac{dn_x}{dt} = 0$. The differential equation for the concentration can then be calculated

$$\frac{dx}{dt} = \frac{d\frac{n_x}{V}}{dt} = \frac{V\frac{dn_x}{dt} - n_x\frac{dV}{dt}}{V^2} = 0 - \frac{n_x}{V}\,\frac{1}{V}\frac{dV}{dt} = -x\mu.$$

The concentration of $X$ thus drops with a rate proportional to itself and $\mu$; this is what we call dilution by growth. This dilution should be matched by a net synthesis of $X$ to maintain a steady state.

#### 6 Deriving the essential self-fabricator relations

In the main text we introduced the essential ingredients for the self-fabricator models: metabolites (with concentrations $\boldsymbol{x}$ and possibly including macromolecules such as lipids or polynucleotides), enzymes (with concentrations $\boldsymbol{e}$), and the ribosome (with concentration $r$). The following set of relations must hold:

$$v_i = e_i k_{\text{cat},i}, \tag{19}$$

$$v_{\text{synth},j} = r k_{\text{cat,rib}} \alpha_j, \tag{20}$$

$$v_{\text{synth,rib}} = r k_{\text{cat,rib}} \alpha_{\text{rib}}, \tag{21}$$

where $v_i$ are the usual metabolic reaction rates, and $v_{\text{synth},j}$ denotes the synthesis rate of enzyme $j$. The factor $\alpha_j$ is the fraction of the ribosome that is allocated to the synthesis of enzyme $j$. It is further assumed that the concentrations of macromolecules add up to a fixed density:

$$\sum_j \rho_j e_j + \rho_{\text{rib}} r = 1, \tag{22}$$

where the $\rho_j$ are volumetric parameters. This density is sometimes modeled as an upper bound instead of a strict equality. In SI7 we show that this is mathematically equivalent when growth is maximized. Imposing the steady state assumption on all metabolites gives a first set of relations between the fluxes:

$$\begin{bmatrix} N & -M_{\text{enz}} & -M_{\text{rib}} \end{bmatrix} \cdot \begin{bmatrix} \boldsymbol{v} \\ \boldsymbol{v}_{\text{synth}} \\ v_{\text{synth,rib}} \end{bmatrix} = \mu \boldsymbol{x}, \tag{23}$$

where $M_{\text{enz}}$ and $M_{\text{rib}}$ are the stoichiometric matrices that denote which metabolites are consumed during the synthesis of enzymes and the ribosome. The steady state assumption for the enzymes and ribosome yield

$$e_j = \frac{v_{\text{synth},j}}{\mu}, \qquad r = \frac{v_{\text{synth,rib}}}{\mu}, \tag{24}$$

because we assume that the concentrations of these compounds solely decline by dilution. Combining this with Equation (19) gives a second relation for the fluxes

$$v_{\text{synth},j} = \mu \frac{v_j}{k_{\text{cat},j}}. \tag{25}$$

To get a relation with the synthesis rate of the ribosome, we can use Equations (20) and (21) to get

$$r = \frac{1}{k_{\text{cat,rib}} \sum_j \alpha_j} \left( \sum_j v_{\text{synth},j} + v_{\text{synth,rib}} \right) = \frac{1}{k_{\text{cat,rib}}} \left( \sum_j v_{\text{synth},j} + v_{\text{synth,rib}} \right),$$

where the last equality is true because the $\alpha_j$-variables are fractions, so that $\sum_j \alpha_j = 1$. Combining this relation with Equation (22) gives the third relation between the fluxes in the model

$$v_{\text{synth,rib}} = \mu \frac{\sum_j v_{\text{synth},j} + v_{\text{synth,rib}}}{k_{\text{cat,rib}}}, \tag{26}$$

Then, using Equation (24) for the last time, we see that in combination with (22) it gives the fourth relation

$$\sum_j \rho_j v_{\text{synth},j} + \rho_{\text{rib}} v_{\text{synth,rib}} = \mu. \tag{27}$$

If we put all the derived relations together, we get

$$\begin{aligned} \underset{\boldsymbol{v}, \boldsymbol{v}_{\text{synth}}}{\text{maximize}} \quad & \mu \\ \text{subject to} \quad & \begin{bmatrix} N & -M_{\text{enz}} & -M_{\text{rib}} \end{bmatrix} \cdot \begin{bmatrix} \boldsymbol{v} \\ \boldsymbol{v}_{\text{synth}} \\ v_{\text{synth,rib}} \end{bmatrix} = \mu \boldsymbol{x} \\ & v_{\text{synth},j} = \mu \frac{v_j}{k_{\text{cat},j}} \\ & v_{\text{synth,rib}} = \mu \frac{\sum_j v_{\text{synth},j} + v_{\text{synth,rib}}}{k_{\text{cat,rib}}} \\ & \sum_j \rho_j v_{\text{synth},j} + \rho_{\text{rib}} v_{\text{synth,rib}} = \mu. \end{aligned} \tag{28}$$

Or, in matrix form:

$$\begin{bmatrix} N & -M_{\text{enz}} & -M_{\text{rib}} \\ \left[\frac{\mu}{k_{\text{cat},j}}\right]_{r \times r} & -\mathbb{I}_{r \times r} & \mathbf{0}_{r \times 1} \\ \mathbf{0}_{1 \times r} & \mathbf{1}_{1 \times r} & 1 - \frac{k_{\text{cat,rib}}}{\mu} \\ \mathbf{0}_{1 \times r} & [\rho_j]_{1 \times r} & \rho_{\text{rib}} \end{bmatrix} \cdot \begin{bmatrix} \boldsymbol{v} \\ \boldsymbol{v}_{\text{synth}} \\ v_{\text{synth,rib}} \end{bmatrix} = \begin{bmatrix} \mu \boldsymbol{x} \\ \mathbf{0}_{r \times 1} \\ 0 \\ \mu \end{bmatrix} \tag{29}$$

where $\left[\frac{\mu}{k_{\text{cat},j}}\right]_{r \times r}$ denotes the diagonal matrix for which the $j^{\text{th}}$ diagonal element is $\frac{\mu}{k_{\text{cat},j}}$. The row vector $[\rho_j]_{1 \times r}$ contains the volumetric constants $\rho_j$. We can rewrite these equalities as homogeneous constraints by realizing that $\mu$ is a variable

$$A(\boldsymbol{x},\mu)\cdot\begin{bmatrix}\boldsymbol{v}\\ \boldsymbol{v}_{\text{synth}}\\ v_{\text{synth,rib}}\\ \mu\end{bmatrix}=\begin{bmatrix}N & -M_{\text{enz}} & -M_{\text{rib}} & -\boldsymbol{x}\\ \left[\frac{\mu}{k_{\text{cat},j}}\right]_{r\times r} & -\mathbb{I}_{r\times r} & \mathbf{0}_{r\times 1} & 0\\ \mathbf{0}_{1\times r} & \mathbf{1}_{1\times r} & 1-\frac{k_{\text{cat,rib}}}{\mu} & 0\\ \mathbf{0}_{1\times r} & [\rho_j]_{1\times r} & \rho_{\text{rib}} & -1\end{bmatrix}\cdot\begin{bmatrix}\boldsymbol{v}\\ \boldsymbol{v}_{\text{synth}}\\ v_{\text{synth,rib}}\\ \mu\end{bmatrix}=\mathbf{0}. \tag{30}$$

This is the form that we proceed with in the main text.

#### 7 Four different ways to deal with an unrealistically low cell density

We presented the density constraint in self-fabrication models as an equality instead of an inequality:

$$\sum_j \rho_j e_j + \rho_{\text{rib}} r = 1.$$

This implies that the maximal density must be met. In [3] we show that this equality is a consequence of assuming that the volume of a cell, $V$, is the sum of the volume of its components. We denote by $n_{x_i}, n_j, n_r$ the copy numbers (in moles) of metabolite $i$, enzyme $j$ and the ribosome, and by $\rho_i, \rho_j, \rho_{\text{rib}}$ the volume occupied by a mole of the corresponding compound. We then get

$$\sum_i \rho_i n_{x_i} + \sum_j \rho_j n_j + \rho_{\text{rib}} n_r = V,$$

where the first sum, the volume contribution of metabolites, is often neglected. If we indeed ignore this contribution, and divide by the volume on both sides, we get

$$\sum_j \rho_j e_j + \rho_{\text{rib}} r = 1. \tag{31}$$

The reason that this equality can not be an inequality is thus that the volume would then not be properly defined anymore.

However, the self-fabrication models that we reviewed all have a constraint which is related to nutrient uptake: either a limited membrane area [7, 8] or just a limited uptake rate [9]. At so-called nutrient-limited conditions, this constraint causes the slow inflow of nutrients. In such a situation, one could imagine that the further processing of this inflow of nutrients does not take up many enzymes. The need for cytosolic enzymes would be less than what fits in the cytosol, and Equation (31) would not be satisfied.

Such an excess of cellular volume gives rise to unrealistic artifacts in optimization models. For example, if no appropriate measures are taken in [9], the excess volume is filled up with an unrealistically high fraction of RNA (because this is metabolically cheaper to produce than excess protein). To prevent this from happening, four different approaches have been used.

In their ME-model, O'Brien et al. [9] model a global saturation factor of enzymes that we will call $\bar{f}$. This saturation factor determines the effective catalytic rate of metabolic enzymes in comparison to the maximal catalytic rate,

$$\bar{f} = \frac{k_{\text{eff}}}{k_{\text{cat}}},$$

and thereby forms a zero order approximation to enzyme kinetics. In their optimization procedure, the authors first assume that all proteins work at their maximal rate. They maximize the growth rate, while demanding that any excess volume is filled up with a certain "dummy protein". If at maximal growth rate the dummy protein is indeed produced, they fix that growth rate and minimize the global saturation factor $\bar{f}$. When this saturation factor is decreased, the need for enzymes is increased, and therefore the excess volume in the cell will vanish. In other words, the authors assume that the cell will fill up any excess volume with the same proteins that it is already using, but that these proteins will now be less efficiently used.

Another option, used by [10,11] is to just consider the density constraint in (31) as an inequality: $\sum_j \rho_j e_j + \rho_{\text{rib}} r \leq 1$. This indeed solves the problem of having excess volume, but an open question remains: what gave rise to the volume of the cell? These models are thus based on the assumption that there is some other, non-modeled, cause that determines the volume of the cell. This volume then gives an upper bound to the cellular content, but the cell can also be only half-full.

Molenaar et al. [8] took a different approach by modeling a cellular shape factor, with which the shape and size of cells can be adjusted. As such there will never be excess volume, since the cell could increase its surface-to-volume ratio by becoming smaller. Hence, the inflow of nutrients can be balanced to the cytosolic capacity for processing these nutrients. This mechanism makes sure that the two constraints always coincide, so that, in accordance with the described extremum principle [3, 12], the gradual switch of overflow metabolism cannot be modeled.

As a last alternative, in [3] we assumed that the ribosome is not always fully occupied. In the main text we introduced the ribosome allocation fractions, $\alpha_i$, that capture which fraction of the ribosome is currently translating enzyme $i$. We then stated that these fractions should sum up to 1. However, during the actual optimization we use

$$\sum_j \alpha_j \leq 1.$$

We thus essentially assume that the excess volume of the cell is filled up with ribosomes that are not occupied. This approach is comparable to the approach of [9]: O'Brien et al. model all metabolic proteins to be unsaturated, while we concentrated this lower saturation on the ribosome.

As a last comment, the above artifacts of having excess volume will be far more abundant in non-kinetic models. This is because, through kinetic effects, additional proteins can almost always be used to get a (small) growth rate benefit. For example, in [3], we include product inhibition. In nutrient-limited conditions, the optimal solution is to express cytosolic proteins to a high level. This will keep all metabolite concentrations in the cell low, because the inflow of nutrients is immediately processed. This will alleviate some of the product inhibition for the transporters, and will therefore lead to faster growth. See [12] for a more elaborate analysis of this scenario.

#### 8 Metabolism and Expression models still fit the standard form

In SI6 we derived the following matrix equation that forms the essential set of equations for self-fabrication models:

$$A(\boldsymbol{x},\mu)\cdot\begin{bmatrix}\boldsymbol{v}\\ \boldsymbol{v}_{\text{synth}}\\ v_{\text{synth,rib}}\\ \mu\end{bmatrix}=\begin{bmatrix}N & -M_{\text{enz}} & -M_{\text{rib}} & -\boldsymbol{x}\\ \left[\frac{\mu}{k_{\text{cat},j}}\right]_{r\times r} & -\mathbb{I}_{r\times r} & \boldsymbol{0}_{r\times 1} & 0\\ \boldsymbol{0}_{1\times r} & \boldsymbol{1}_{1\times r} & 1-\frac{k_{\text{cat,rib}}}{\mu} & 0\\ \boldsymbol{0}_{1\times r} & [\rho_j]_{1\times r} & \rho_{\text{rib}} & -1\end{bmatrix}\cdot\begin{bmatrix}\boldsymbol{v}\\ \boldsymbol{v}_{\text{synth}}\\ v_{\text{synth,rib}}\\ \mu\end{bmatrix}=\boldsymbol{0}. \tag{32}$$

In Metabolism and Expression models, more cellular components are included. For example, in O'Brien et al. [9], an RNA-polymerase, mRNAs, and tRNAs are added. To these components a synthesis and a dilution rate are associated, which should be balanced. The dilution rate is, as with all cellular compounds, proportional to the growth rate and to the concentration of the compound. The concentration of the compound can be related to the necessary flux of the catalyzed reaction via the catalytic rate of the catalyst.

For example, RNA-polymerases catalyze transcription of RNA. It is assumed that the rate of transcription is proportional to the length of the RNA-molecule, measured by the number of nucleotides. The total flux that needs to be catalyzed is the sum of the transcription rates of all RNA-molecules multiplied by their number of nucleotides

$$v_{\text{total transcription}}=\sum_i v_{\text{transcription},RNA_i}\text{length}(RNA_i).$$

The RNA-polymerases will have a certain catalytic rate, so that

$$v_{\text{total transcription}}=k_{\text{cat,RNAP}}rnap,$$

where $rnap$ is the concentration of RNA-polymerase. Combining these equations gives

$$rnap=\frac{1}{k_{\text{cat,RNAP}}}\sum_i v_{\text{transcription},RNA_i}\cdot\text{length}(RNA_i).$$

We can now calculate the dilution rate by using this in $v_{\text{dilution},RNAP}=\mu\cdot rnap$. The synthesis rate of RNAP should equal this dilution term,

$$v_{\text{synthesis},RNAP}=\frac{\mu}{k_{\text{cat,RNAP}}}\sum_i v_{\text{transcription},RNA_i}\cdot\text{length}(RNA_i). \tag{33}$$

This derivation is done in a similar fashion for the synthesis rates of RNAs.

An additional complexity that is added in ME models is that the catalytic rates of some catalysts, such as the ribosome and RNA-polymerase, are no longer assumed to be constant. Rather, a nonlinear, growth rate dependent catalytic rate is inferred from experimental data. Equation (33) thus becomes

$$v_{\text{synthesis},RNAP} = \frac{\mu}{k_{\text{cat,RNAP}}(\mu)} \sum_i v_{\text{transcription},RNA_i} \cdot \text{length}(RNA_i). \quad (34)$$

This relation is still linear in the reaction rates and nonlinear in the growth rate. Therefore, we can extend Equation (32) to incorporate this constraint:

$$\begin{bmatrix} N & -M_{\text{enz}} & -M_{\text{rib}} & \mathbf{0}_{1\times q} & 0 & -\boldsymbol{x} \\ \left[\frac{\mu}{k_{\text{cat},j}}\right]_{r\times r} & -\mathbb{I}_{r\times r} & \mathbf{0}_{r\times 1} & \mathbf{0}_{1\times q} & 0 & 0 \\ \mathbf{0}_{1\times r} & \mathbf{1}_{1\times r} & 1-\frac{k_{\text{cat,rib}}}{\mu} & \mathbf{0}_{1\times q} & 0 & 0 \\ \mathbf{0}_{1\times r} & \mathbf{0}_{1\times r} & 0 & \left[\frac{\mu\cdot\text{length}(RNA_i)}{k_{\text{cat,RNAP}}(\mu)}\right]_{1\times q} & -1 & 0 \\ \mathbf{0}_{1\times r} & [\rho_j]_{1\times r} & \rho_{\text{rib}} & [\rho_{\text{j,RNA}}]_{1\times q} & \rho_{\text{RNAP}} & -1 \end{bmatrix} \cdot \begin{bmatrix} \boldsymbol{v} \\ \boldsymbol{v}_{\text{synth}} \\ v_{\text{synth,rib}} \\ \boldsymbol{v}_{\text{synth,RNA}} \\ v_{\text{synth,RNAP}} \\ \mu \end{bmatrix} = \mathbf{0}. \quad (35)$$

Here, we have used $q$ to denote the number of different RNAs that should be produced. The relations between the synthesis rates of RNAs and the synthesis rates of enzymes can be added as well, but is here, for conciseness, omitted. This Equation still fits the general form

$$A(\boldsymbol{x},\mu) \cdot \begin{bmatrix} \boldsymbol{v} \\ \mu \end{bmatrix} = \mathbf{0}.$$

Concluding, the Metabolism and Expression models add variables and constraints as compared to more basic self-fabrication models, but these extensions do not change the form of the mathematical optimization problem.
